# FoxO factors preserve airway epithelial homeostasis by coordinating adaptive stress responses

**DOI:** 10.1101/2024.01.31.578231

**Authors:** Karin Uliczka, Judith Bossen, Ulrich M. Zissler, Christine Fink, Xiao Niu, Mario Pieper, Ruben D. Prange, Christina Vock, Christina Wagner, Mirjam Knop, Ahmed Abdelsadik, Sören Franzenburg, Iris Bruchhaus, Michael Wegmann, Carsten B. Schmidt-Weber, Peter König, Petra Pfefferle, Holger Heine, Thomas Roeder

## Abstract

Airway epithelia must maintain functional and structural homeostasis despite exposure to diverse environmental stressors. FoxO transcription factors are well suited to this task, acting as integration hubs that translate extrinsic and intrinsic signals into appropriate physiological responses. We show that FoxO factors in *Drosophila*, mouse, and human airway epithelial cells (AECs) undergo nuclear translocation in response to stressors including hypoxia, temperature, and oxidative stress. In human airway epithelial cell lines, individual hFOXO factors show distinct, cell-type-specific activation patterns. In *Drosophila*, hypoxia was the only stressor among those tested that triggered a dfoxo-dependent innate immune response. Because *Drosophila* possesses a single FoxO ortholog, loss-of-function analysis directly demonstrated that dfoxo is required for stress resistance in the airway epithelium. Reduced FoxO expression was similarly observed in mouse models of asthma and in airway samples from asthma patients, paralleling the increased stress sensitivity seen upon FoxO loss in flies. While our data do not establish that reduced FOXO levels cause asthma, they indicate that FoxO-dependent stress pathways are altered in diseased airways across species. Together, these findings support a model in which FoxO coordinates adaptive epithelial stress responses that maintain airway homeostasis under environmental challenge.

## 1 INTRODUCTION

Airway epithelial cells (AECs) form the barrier between the respiratory organs and the environment. Therefore, AECs are permanently exposed to physical stressors, pathogens, environmental pollutants, and allergens. A significant task of AECs is to orchestrate the lung’s responses to these environmental influences, thereby maintaining airway structural integrity and immune homeostasis(Hewitt & Lloyd, 2021; Weisberg *et al*, 2021; Whitsett & Alenghat, 2015). Impairment of these functions is associated with developing chronic lung diseases such as asthma, chronic obstructive pulmonary disease (COPD), or pulmonary fibrosis, supporting the hypothesis that these diseases are primarily epithelial(Celebi Sozener *et al*, 2020; Heijink *et al*, 2020; Hellings & Steelant, 2020; Lambrecht & Hammad, 2012). This interpretation is further supported by the fact that carriers of risk alleles for these diseases are particularly prone to develop deregulated epithelial immune functions (Hellings & Steelant, 2020; Pividori *et al*, 2019; Raby *et al*, 2023) and inappropriate repair responses (Burgoyne *et al*, 2021; Chung & Adcock, 2008). Significant disease hallmarks comprise airway remodeling and prolonged, persistent inflammatory reactions (McAlinden *et al*, 2019; Proud & Leigh, 2011; Rodriguez-Colman *et al*, 2024; Spann *et al*, 2014; Zepp & Morrisey, 2019).

This central property of AECs to respond appropriately to a wide variety of intrinsic and extrinsic signals requires signaling systems that react to these signals and connect their activation with an appropriate cellular response. The transcription factors of the FoxO family precisely fulfill these requirements (Gui & Burgering, 2022; Orea-Soufi et al, 2022). They are critically involved in various cellular processes, including apoptosis, cell survival, oxidative stress responses, and energy metabolism (Graves & Milovanova, 2019). While the fruit fly *Drosophila* melanogaster has only one FoxO gene (dfoxo) (Junger et al, 2003), mammals have four FoxO factors (in humans: hFOXO1, hFOXO3, hFOXO4, and hFOXO6). In all these organisms, these factors act as integration hubs for different cellular signaling systems, such as the insulin and JNK pathways, responding to various signals (Cheng, 2019; Greer & Brunet, 2005; Santos et al, 2023; van der Vos & Coffer, 2011; Wagner et al, 2021).

The interest in FoxO factors is largely based on research efforts in two different scientific fields: aging and research on a broad spectrum of chronic diseases. In aging studies, increased FoxO activity usually correlates with a prolonged lifespan. This correlation, initially found in models such as C. elegans and *Drosophila* (Giannakou et al, 2007; Sun et al, 2017), is further supported by the fact that hFOXO3 is one of the most consistently replicated longevity genes (Donlon et al, 2022; Flachsbart et al, 2009). On the other hand, deregulation of hFOXO activities has been associated with various chronic human diseases(Manolopoulos et al, 2010; Orea-Soufi et al., 2022; van der Horst & Burgering, 2007; Xing et al, 2018). This causal relationship between altered hFOXO activity and disease pathogenesis has also been demonstrated for chronic lung diseases. Thus, a deficiency of the transcription factor hFOXO1 contributes to developing pulmonary hypertension(Savai et al, 2014; Selle et al, 2022). Deficiencies in *hFOXO3*, on the other hand, correlate with COPD and idiopathic pulmonary fibrosis (IPF)(Al-Tamari et al, 2018; Hwang et al, 2011). Especially, the correlation with IPF has recently been supported in various studies (Nho et al, 2011; Nho et al, 2013; Xin et al, 2018). Moreover, alterations in the expression of *hFOXO* factors were associated with various cancer entities including lung cancer(Hornsveld et al, 2018; Jiramongkol & Lam, 2020; Mikse et al, 2010). In human respiratory epithelial cells, increased activation of the hFOXO signaling pathway directly controls an epithelial immune response, a mechanism first demonstrated in *Drosophila*(Becker et al, 2010; Seiler et al, 2013). Consistently, rhinoviral infection activates hFOXO3 in airway epithelia and helps to orchestrate antiviral responses(Gimenes-Junior et al, 2019). Recently, general aspects of FoxO biology in epithelial tissues were elucidated in *Drosophila* (Fink et al, 2016; Wagner et al., 2021).

Despite extensive evidence linking FOXO factors to immunity, tissue repair, metabolism, and chronic lung diseases, their role as central regulators of airway epithelial stress adaptation remains poorly understood. Airway epithelial cells are continuously exposed to a broad spectrum of environmental and endogenous stressors, yet it is unclear whether FOXO signaling constitutes a conserved stress-response system that integrates these diverse stimuli and promotes epithelial resilience. Furthermore, while altered FOXO expression has been reported in several chronic lung diseases, it remains unknown whether impaired FOXO-dependent stress responses contribute to the increased vulnerability of diseased airways to environmental challenges.

To gain deeper insight into the role of FoxO-mediated signaling in airway epithelial cells (AECs), we performed complementary analyses in *Drosophila*, mice, and humans. Our studies revealed that FoxO factors activate in AECs in response to diverse environmental stressors across species. However, the downstream responses are likely species-specific. In human cells, nuclear translocation of hFOXO factors varied depending on the cell type and hFOXO isoform. Furthermore, dfoxo deficiency reduced stress tolerance in *Drosophila*, showing a functional requirement for FoxO in epithelial adaptation to environmental stress. Consistent with these findings, mFoxo transcript levels were significantly reduced in mouse models of asthma. hFOXO expression was also decreased in airway samples from asthma patients, supporting the relevance of FoxO-associated stress response pathways in chronic airway disease.

## 2 RESULTS

### 2.1 Hypoxia induces a dfoxo-dependent immune response in the *Drosophila* airways

Hypoxia is a vital airborne stressor that can induce a variety of responses. We have used the fruit fly *Drosophila* as a model to study these responses. To investigate which organ responds most sensitively to a hypoxic episode, we used the LDH-Gal4/UAS-nGFP reporter strain, which shows nuclear expression of GFP under transcriptional control of the murine LDH-A enhancer when the animals were subjected to hypoxia (Lavista-Llanos et al, 2002). Hypoxia elicited a highly reproducible response of this reporter in the airway epithelium but not in other organs (Figure 1A; background staining of the intestine represents autofluorescence caused by ingested food). This result suggests that the airway epithelium is strongly affected by hypoxia and more susceptible to this stressor than other organs. In preliminary experiments, we observed that the epithelial immune system is susceptible to environmental cues. Therefore, we performed a transcript analysis of isolated tracheae, focusing on selected antimicrobial peptides (AMPs) (Kallsen et al, 2015; Wagner et al, 2008; Wagner et al, 2009). We exposed *Drosophila* larvae to mild hypoxia (5 % O_2_ for 2 h) and found that some of the genes studied, namely cecropin, attacin, and drosomycin, were consistently and significantly up-regulated (p<0.05; Figure. 1B).

**Figure 1:**
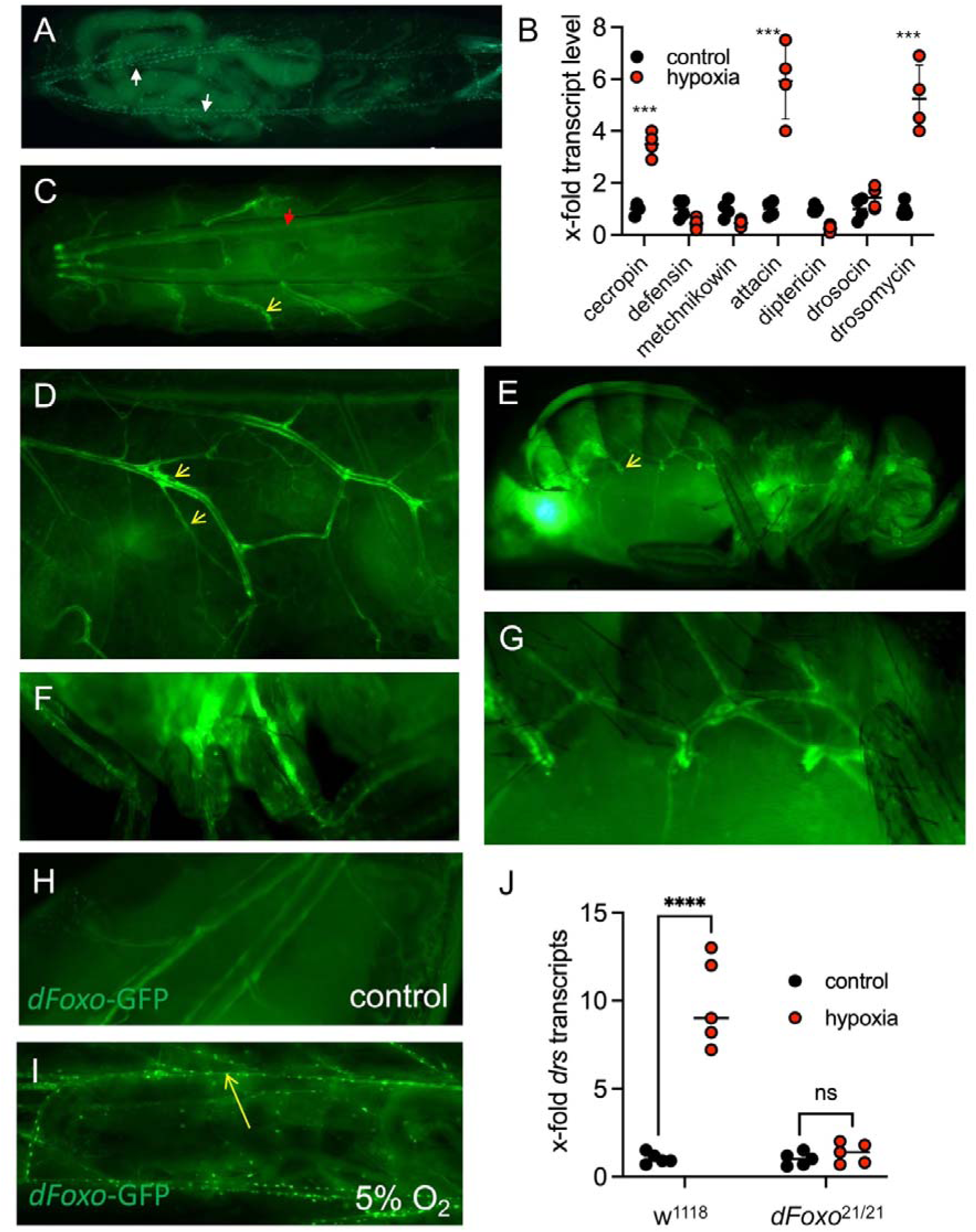
Hypoxia induces a dfoxo-dependent activation of the immune response in *Drosophila* airway epithelia. A) Early third-instar larvae of the LDH-Gal4/UAS-nGFP genotype were subjected to a brief period of mild hypoxia (2 h, 5% oxygen). Nuclear expression of GFP was observed only in the airway epithelium (arrows). Background staining of the intestine is due to the autofluorescence of ingested material. B) qRT-PCR analysis of the transcript levels of the major antimicrobial peptide (AMP) gene levels in the trachea of larval *Drosophila* in response to a hypoxic episode. C, D) Hypoxia treatment of a drosomycin reporter strain (Drs-GFP) induced GFP expression only in the airway epithelium. C) In the overview, expression is restricted to the primary and secondary branches (yellow arrows), whereas the main trunks show no or little expression (red arrows). D) At higher magnification, expression in the primary and secondary branches is more evident. E-G) Using the same line, hypoxia-induced expression of the drosomycin reporter was also observed in the adult tracheal system. E) At low magnification, the peripheral tracheal system is visible through the cuticle (yellow arrows). F) At higher magnification, tracheal structures directly adjacent to the spiracular opening and G) within the legs are visible. dfoxo-gfp targeting the tracheal system under H) control and I ) hypoxic conditions. Nuclear translocation is seen in I as the dot-like structure. J) Hypoxia-induced transcript levels of drosomycin in larval trachea of control (left) and *dfoxo*-deficient animals. Mean values ± S.D. ≥ 3, ** p<0.01.

We then used the drosomycin-GFP (Drs-GFP) reporter line to elucidate the spatial dimension of this antimicrobial response reaction to hypoxia. Here, the expression was strongly increased by hypoxic episodes (Figure 1C), and the expression appeared first in the most distal part of the airway system, where it was observed in most treated animals (yellow arrows; Figure 1C, D). Main branches showed little or no response under these conditions, whereas secondary and terminal branches showed strong fluorescence (red arrows; Figure. 1C, D). After prolonged hypoxia, all major branches (except the dorsal trunks) showed robust expression of this antimicrobial (antifungal) peptide gene. In contrast, other organ systems showed no induction of GFP expression in response to hypoxia (Figure. 1C, D). When adult animals were exposed to hypoxia, strong GFP expression was observed throughout the tracheal system (white arrows, Figure 1E). Detailed analysis revealed strong expression in tracheae close to the surface of the animal, namely those in the legs (Figure. 1F) and those regions immediately adjacent to the tracheal opening (Figure 1G).

To identify the underlying mechanism of this antimicrobial response, we targeted XFP-tagged transcription factors known to be involved in the regulation of antimicrobial responses and analyses their nuclear translocation as a proxy for their activation. Flies expressing XFP-tagged transcription factors [UAS-relish-yfp, UAS-dorsal-gfp, UAS-dif-cfp (Bettencourt et al, 2004), and UAS-dfoxo-gfp] were targeted to the airways by crossing with ppk4-Gal4. We used the ppk4-Gal4 driver because it shows more pronounced expression in later L3 larval stages than the btl-Gal4 driver, which was the developmental stage at which we performed these experiments (Liu et al, 2003; Wagner et al., 2008). These animals were exposed to hypoxia. Neither Relish nor Dif or Dorsal showed any nuclear translocation following hypoxia. Under normoxic conditions, dfoxo-GFP localizes to the cytoplasm of airway epithelial cells (Figure 1H). In response to hypoxia, an almost complete translocation of dFoxo into the nucleus was observed (Figure 1I).

Based on the above results, we tested whether dfoxo is necessary for hypoxia-induced expression of drosomycin in the trachea. In dFoxo-deficient animals (dfoxo^21/21^), hypoxia-induced drosomycin expression was absent, demonstrating that dfoxo is essential for this immune response (Figure 1J). The observation that not al l AMPs are regulated by dfoxo is in accordance with earlier studies in the flies’ intestine (Fink *et al*., 2016).

### 2.2 dfoxo translocation in response to different types of stimuli

Having shown that hypoxia is a major stressor capable of inducing dfoxo translocation in the trachea, we investigated to what extent other stressors also induce this response. In parallel, we tested whether the other immune-related transcription factors, Dorsal and Relish, respond with nuclear translocation. Ectopically expressed transcription factors labeled with GFP or YFP were introduced into the respiratory tract of *Drosophila* larvae using the Gal4/UAS system (Bettencourt *et al*., 2004). In the untreated control group, Dorsal, Relish, and dfoxo were localized in the cytoplasm (Figure 2A, A’, A’’). Starvation, oxidative stress, UV irradiation, cold stress, and heat stress resulted in a rapid translocation of dfoxo to the nucleus (Figure 2B-F). However, nuclear translocation of Relish (Figure 2B’-F’) and Dorsal (Figure 2B’’-F’’) was not observed, except for a partial nuclear translocation of Relish during starvation stress (Figure 2B’).

**Figure 2:**
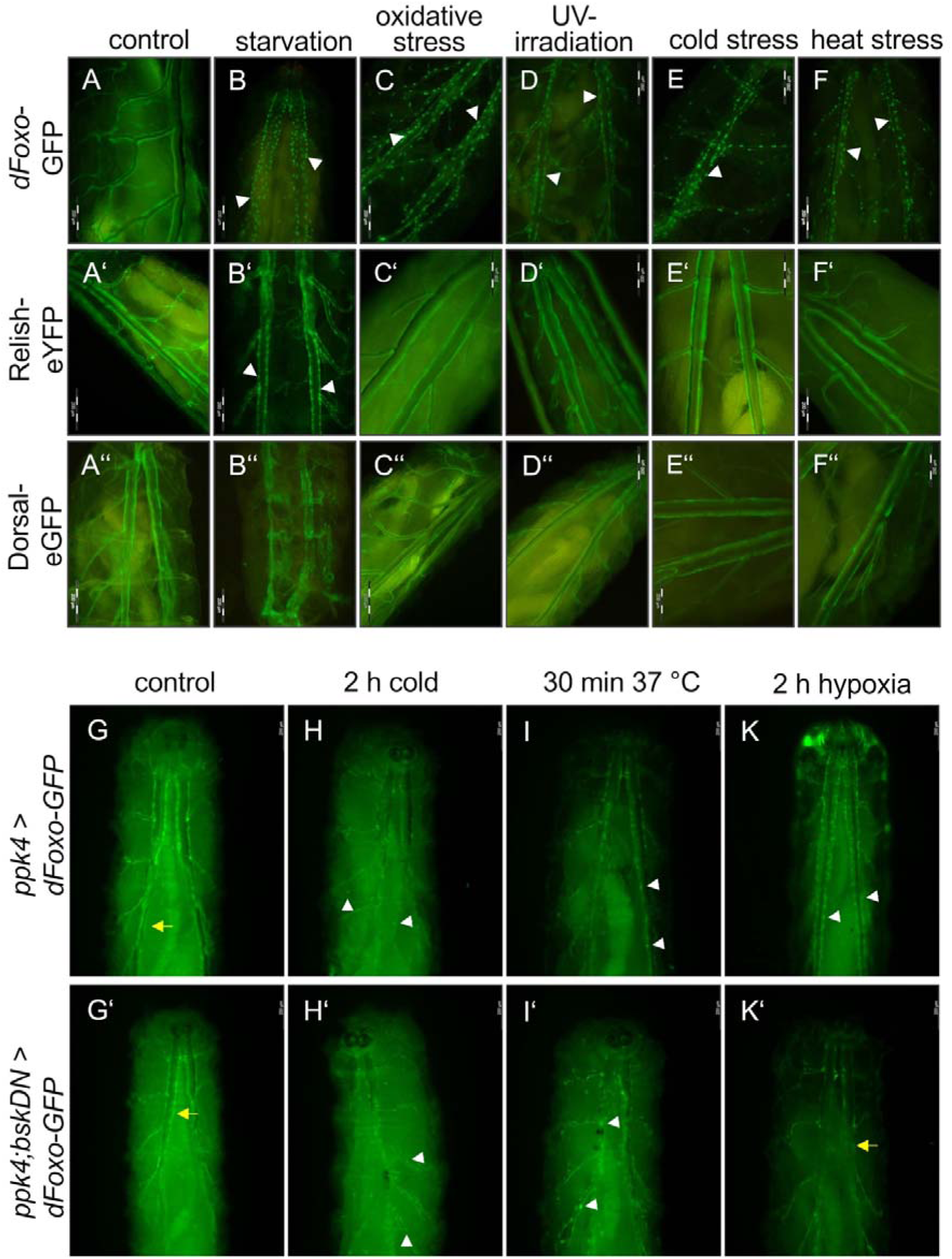
Cellular localizations of the transcription factors dfoxo, relish, and dorsal in the airways of *Drosophila* larvae exposed to different stressors. A-F) *dfoxo-GFP* was expressed in the airways by crossing the driver line *btl*-Gal4 with the corresponding responder line UAS-*dfoxo-gfp*. A) Under non-stressed conditions, fluorescence was restricted to the cytoplasm. B-F) Starvation (B, 12 h of nutritional deprivation), oxidative stress (C, exposure to 100 mM paraquat for 2 h), UV irradiation (D, 254 nm UVC rays), cold stress (E, 2 h at 4°C), and heat stress (F, 30 min at 37°C) induced nuclear translocation of dFoxo-GFP in epithelial cells of the tracheae. A’-F’’) Relish-eYFP and Dorsal-eGFP were expressed by crossing the driver line *btl*-Gal4 with the corresponding responder lines UAS-*relish-eyfp* and UAS-*dorsal-egfp*, respectively. A’, A’’) Under non-stressed conditions, the fluorescence of Relish-eYFP and Dorsal-eGFP was restricted to the cytoplasm. B’, B’’) Except for starvation (B’), the various stressors including C’) oxidative stress, D’) UV irradiation, E’) cold stress, F’) and heat stress did not induce nuclear translocation of Relish-eYFP. B’’-F’’) None of the applied stressors induced nuclear translocation of Dorsal-eGFP. G-K) *ppk4-Gal4XUASdfoxo-gfp* animals remained unstressed (G, control), H) experienced cold, I ) heat, and K) hypoxia. G’-K’) Animals of the genotype *ppk4-Gal4; UAS-bsk^DN^XUAS-dfoxo-gfp* were kept under control conditions. White arrowheads indicate nuclear translocation. Yellow arrows indicate no nuclear translocation. All images were acquired 2 h, or 12 h in the case of starvation after stress stimulation was initiated. Scale bars are 200 µm.

To analyze the underlying activation mechanisms, we also performed these studies when the JNK pathway was knocked down (induced mechanism with Bsk^DN^ by co-expression of the dominant negative JNK allele (Bsk^DN^), Fig. 2G-K’). Under these conditions, the cold-(Fig. 2H, H’) and heat-induced (Fig. 2I, I’) translocation of dfoxo was not dependent on the JNK pathway, whereas hypoxia-induced dfoxo translocation was drastically reduced (Fig. 2K, K’).

### 2.3 Transcriptomic analyses of *Drosophila* airways upon dfoxo deregulation

We performed transcriptome analyses of isolated *Drosophila* larval airways to learn more about the processes involved in deregulated dfoxo expression. To this end, we analyzed the effects of dfoxo deficiency in dfoxo^21/21^ animals (Fig. 3A, B) and those of specific airway-targeted overexpression (Fig. 3C, D). dfoxo deficiency resulted in the upregulation of 768 genes and the downregulation of 1745 genes (>1.5-fold, p<0.05). In contrast, dfoxo overexpression, specifically in the trachea (dfoxo^oe^), resulted in the upregulation of 252 genes and the downregulation of 344 genes. In both experiments, results obtained with control and dfoxo^21/21^or dfoxo^oe^ tracheae were significantly separated in the corresponding PCA plots (Figure 3B, D). Overlaps between the different groups in the Venn diagram were particularly relevant (Figure 3E). We identified 110 genes that were downregulated in dFoxo-deficient tracheae and upregulated in dfoxo^oe^ tracheae. Enriched GO terms for biological processes in this cohort were mainly associated with cuticle development and muscle function and development. In addition, 42 genes were downregulated in dfoxo^oe^flies and upregulated in dfoxo^21/21^flies (Fig. 3E). We also found regulation in the same direction, including those that were downregulated (88 genes) or upregulated (22 genes) in both dfoxo^oe^ and dfoxo^21/21^ flies. All these overlaps were statistically significant (Fisher’s exact test, p<0.05; Fig. 3E). Genes downregulated in dfoxo^21/21^flies were particularly interesting. The following KEGG pathways were enriched: various drug-related metabolic detoxification processes, glutathione metabolism, and amino acid metabolism (Figure 3F). Transcripts that were upregulated in dfoxo^21/21^ airways were associated with the following KEGG pathways: DNA replication, Drug metabolism, and associated terms (Figure 3G). Figure 3H shows those transcripts related to glutathione metabolism that were regulated in dfoxo^21/21^airways, as glutathione metabolism largely overlaps with additional KEGG pathways, including Drug metabolism. In addition, those heat shock proteins (Hsp) that show differential expression (compared to control airways) in dfoxo^21/21^ airways are also listed (Figure 3I).

**Figure 3:**
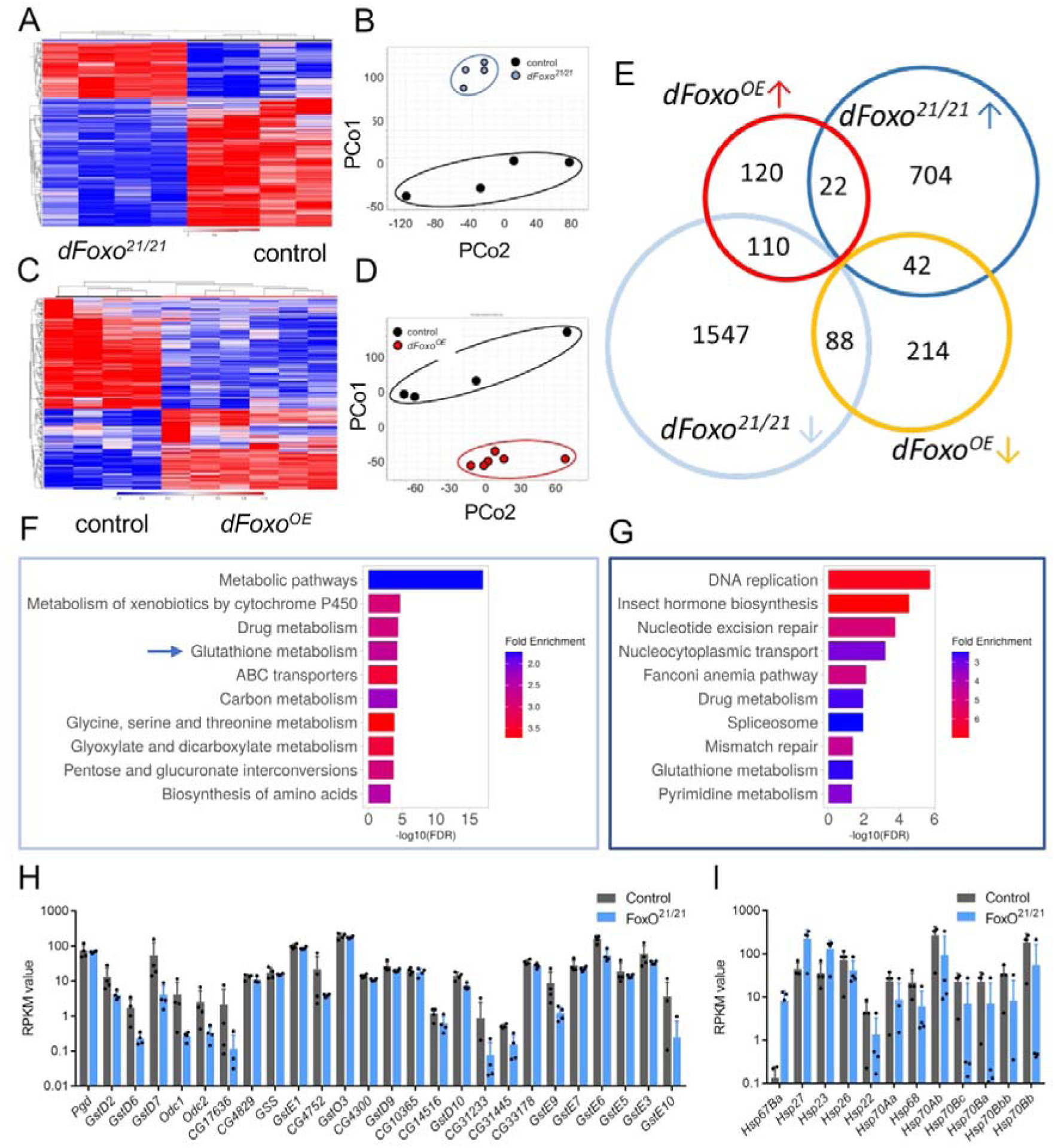
RNAseq analyses of dfoxo deficiency and dfoxo overexpression in the airways of *Drosophila* larvae. Heatmap of differentially expressed genes >1.5 fold, (p<0.05) of airway samples taken from control and *dfoxo^21/21^*animals. B) Red indicates upregulation, blue indicates downregulation. PCoA analysis of the individual biological replicates is shown on the right. C) Heatmap of differentially expressed genes between control airways and those experiencing *dfoxo* overexpression (*dfoxo*^oe^) in the airways. D) PCoA analysis is shown on the right. E) Venn diagram analysis of significantly up- and downregulated genes of comparisons between controls and *dfoxo^21/21^*as well as controls and *dfoxo* overexpression (oe). The significance of overlapping cohorts of genes was calculated with Fisher’s exact test (***p<0.001, *p<0.05). F) Significantly regulated KEGG pathways in the cohort of genes that are downregulated in dfoxo^21/21^. G) Significantly regulated KEGG pathways in the cohort of genes that are upregulated in dfoxo^21/21^ trachea. H) Transcript levels of genes associated with glutathione metabolism, showing both decreased and increased expression, in *dfoxo^21/21^*airways (mean values ± S.D. are shown). I ) Heat-shock protein genes whose transcription is differentially regulated in *dfoxo^21/21^*airways. Genes shown in H and I are significantly different (p< 0.05) from the controls.

Collectively, these transcriptomic analyses identify dFoxo as a central regulator of airway epithelial homeostasis. Loss of dFoxo reduced the expression of genes associated with glutathione-dependent detoxification, xenobiotic metabolism, stress responses, and epithelial maintenance, while increasing transcriptional programs linked to DNA replication, cell-cycle progression, and developmental signaling. Conversely, dfoxo overexpression shifted the transcriptome towards tissue maintenance while suppressing proliferative programs. Together, these reciprocal changes indicate that dFoxo orchestrates coordinated transcriptional remodeling during chronic airway stress. Rather than regulating isolated stress-response genes, dFoxo integrates antioxidant defense, detoxification, innate immune-associated responses, metabolic homeostasis, and epithelial maintenance into a unified adaptive program.

### 2.4 Stress sensitivity of *dfoxo* deficient *Drosophila*

Having shown that various stressors activate dFoxo, we sought to test the hypothesis that dFoxo is essential for an adequate response to stressors. We compared dfoxo^21/21^flies and their control counterparts (yw) after exposure to different stressors (Figure 4). I t is known that daily exposure to cigarette smoke significantly shortens the lifespan of flies (Prange et al, 2018). Our experiment shows that this shortening was greater in dfoxo^21/21^flies than in control flies exposed to the same protocol (9 days for dfoxo^21/21^ flies versus 18 days median lifespan for control fl ies; Figure 4A). A similar reduction was observed when we silenced dfoxo exclusively in the tracheal cells using CRISPR/Cas9 (btl-Gal4; UAS-gRNA-Foxo; UAS-Cas9; Figure 4-supplemental Figure 1). These findings demonstrate that FoxO activity within the airway epithelium itself substantially contributes to epithelial resistance against chronic airborne stress. Drought stress also reduced the mean lifespan of adult flies. This lifespan-shortening effect was enhanced in dfoxo^21/21^ flies (32 h for dfoxo^21/21^ flies versus 46 h for control flies: Figure 4B). Finally, we examined the effects of hypoxia using a hypoxia-induced larval escape assay (Figure 4C) and the hypoxia recovery assay with adult flies (Figure 4D). In the former assay, dfoxo^21/21^ larvae were more sensitive to hypoxia than control larvae because they left the medium earlier (Figure 4C). In the latter assay, the dFoxo^21/21^flies recovered much more slowly than the control flies (Figure 4D). Taken together, these experiments demonstrate the increased stress sensitivity of dfoxo-deficient animals.

**Figure 4:**
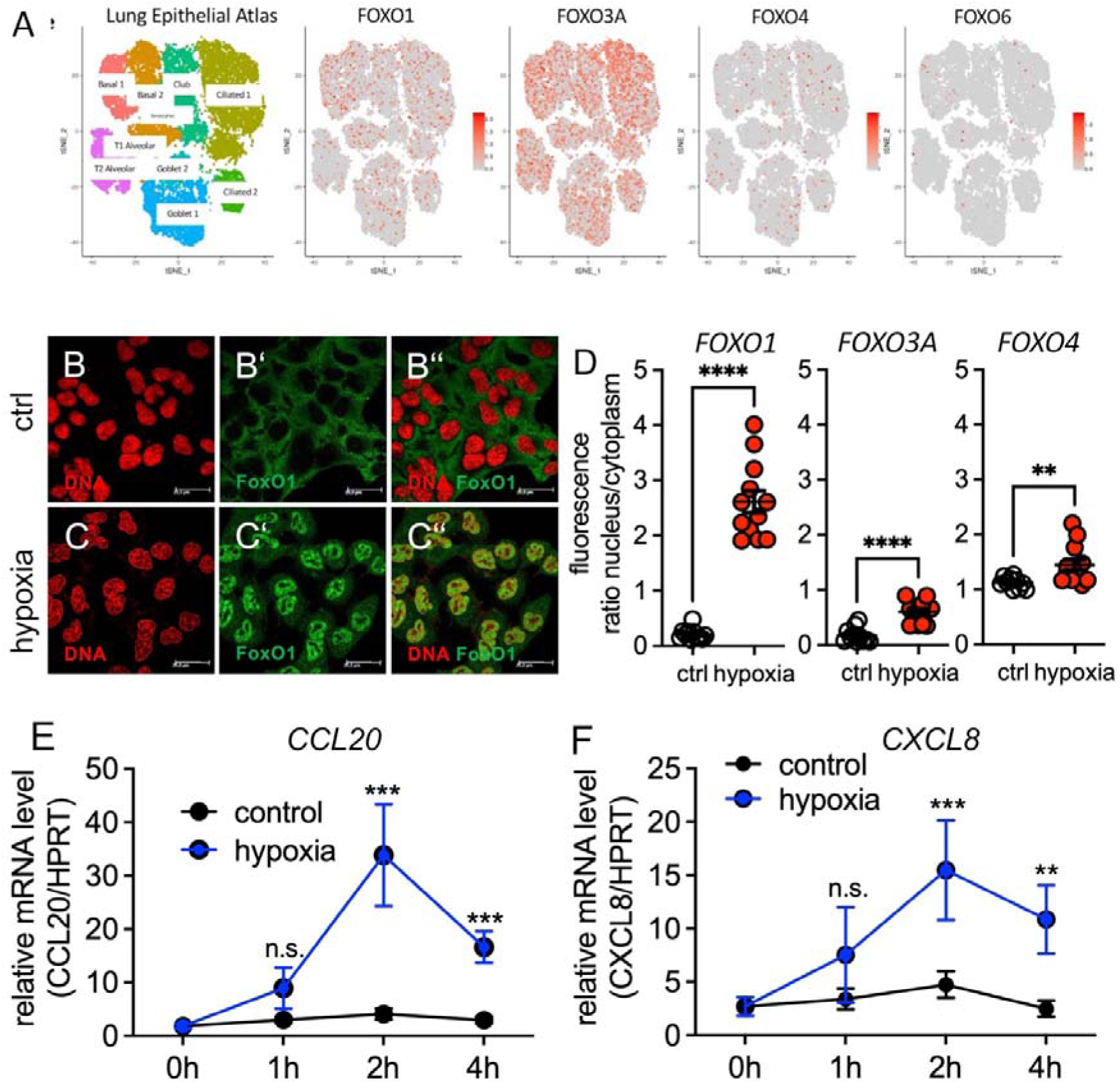
Reduced stress resistance of dfoxo^21/21^flies. A) Survival of *dfoxo*-deficient (*dfoxo^21/21^*) and control adult flies exposed to cigarette smoke daily (n>10). B) Survival of *dfoxo*-deficient and control flies exposed to desiccation. C) Lawn-leaving assays of *Drosophila* 3^rd^ instar larvae exposed to hypoxia. D) *dfoxo*-deficient and control flies were briefly anesthetized with N_2_ and the time taken to full recovery was measured. For A, B, N≥100, for C, D, N=30-110. Shown are mean values ± SEM. * means p<0.05, ** means p<0.01, *** means p<0.005. Survival curves were compared by the log-rank (Mantel–Cox) test.

### 2.5 Effects of hypoxia on hFOXO in human airway epithelial cells

We wondered whether the response to hypoxia observed in *Drosophila* would be similar in the human system. To analyze this, we first investigated whether the different hFOXO factors are expressed in the human lung. We took advantage of the human lung atlas (Vieira Braga et al, 2019) and evaluated the expression of the four hFOXO members (Figure 5A). hFOXO3 and, to a lesser extent, hFOXO1 and hFOXO4 showed broad expression in almost all epithelial cell types (Figure 5A). In contrast, hFOXO6 is expressed at much lower levels (Figure 5A). We used the lung epithelial cell line A549, exposed it to hypoxic conditions, and analyzed the translocation properties of the three major hFOXO factors, hFOXO1, 3, and 4, by immunohistochemistry (Figure 5B). hFOXO1 showed a robust nuclear translocation in response to hypoxia (Figure 5C), whereas the hypoxia-induced nuclear localization of hFOXO3 and hFOXO4 was less pronounced (Figure 5D). We then investigated the extent to which cytokines are also induced by hypoxia in A549 cells. We focused on two cytokines that were shown to react to stressors in airway epithelial cells reproducibly, CCL20 (Figure 5E) and CXCL8 (Figure 5F) (Guesdon et al, 2015; Ovrevik et al, 2009; Wolf & Moser, 2012). Both cytokines were significantly upregulated in response to hypoxia. Together, these findings suggest that FOXO responsiveness to airway-associated stress is not restricted to *Drosophila* but is also observed in mammalian airway epithelial cells. However, whether this reflects conservation of downstream FOXO-regulated transcriptional programs remains unresolved.

**Figure 5:**
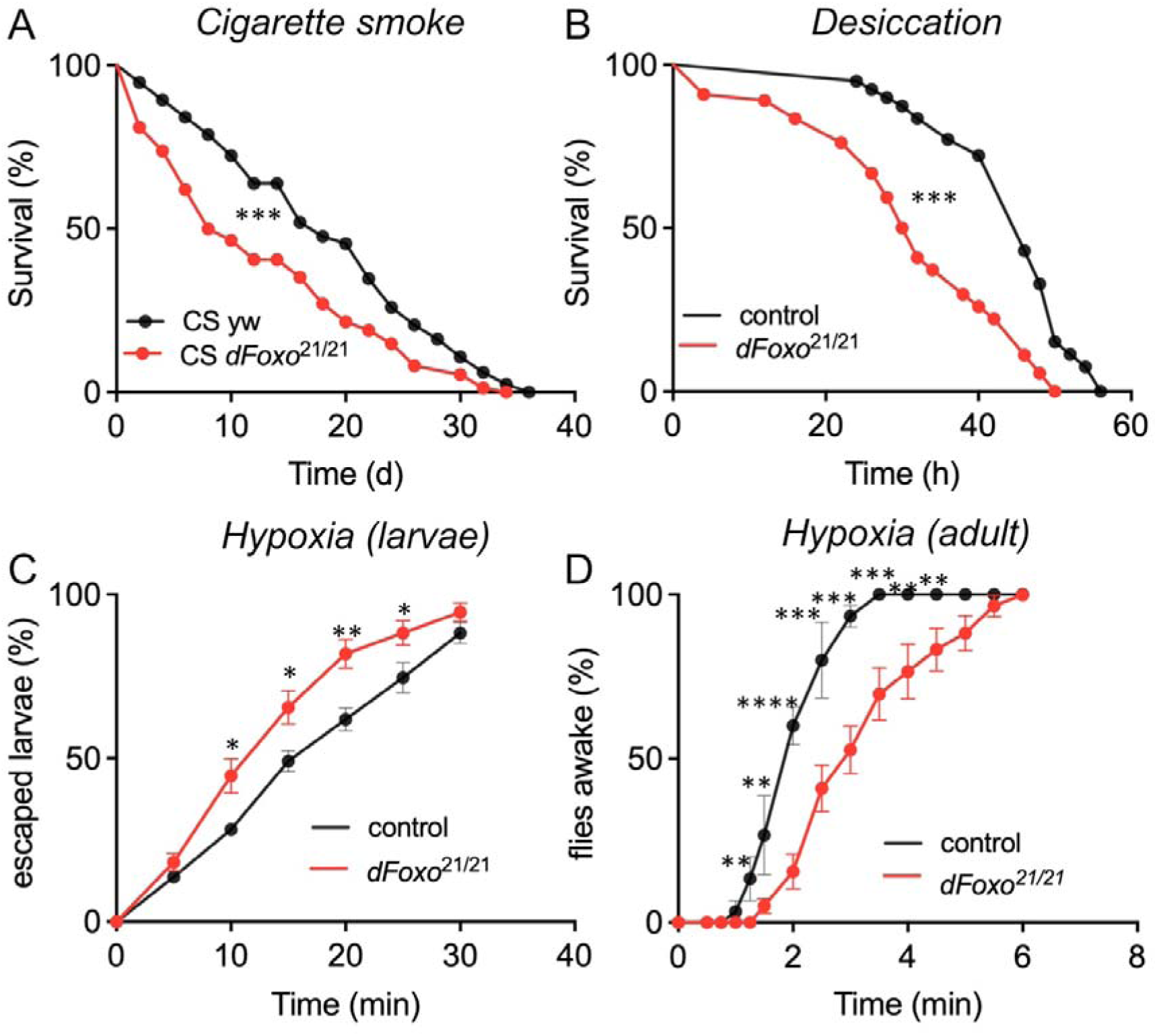
Expression of hFOXO1, hFOXO3, hFOXO4, and hFOXO6 in lung tissues of humans and response to hypoxia. Using the human lung atlas (Vieira Braga *et al*., 2019), expression plots of the four different hFOXO genes were generated (A). Red dots represent positive cells. The assignment to specific epithelial cell types is shown on the left. B-D) Response to hypoxic treatment (5% O_2_) in human A549 cells. B) Control cells show a cytosolic localization of hFOXO1, whereas hypoxia induced an almost complete nuclear translocation (C). (D) Quantitative evaluation of hFOXO translocation in response to hypoxia as indicated by hFOXO1, hFOXO3, and hFOXO4. Each dot represents the mean value obtained for one biological replicate. E, F) Hypoxia-induced expression of cytokines with antimicrobial activities in a time-dependent manner. E) Expression levels of CCL20 in A549 cells treated for different times with hypoxia (5 % O_2_). F) The same type of analysis as in E was performed with CXCL8. Data are presented as the mean and SD (n≥6 per group). Statistical analyses were performed using a one-way ANOVA. *p<0.05, **p<0.01, ***p<0.001.

### 2.6 Stress-induced nuclear translocation of hFOXO factors in human airway epithelial cells

Next, we used human cell lines representing different airway epithelial cell types to evaluate if the stressors identified in the *Drosophila* system also trigger the translocation of hFOXO factors (Figure 6, Figure 6-supplemental Figure 1A-J). From many different human airway epithelial cell lines, we selected two lines that differ as much as possible in their properties and origins and thus represent a large part of the diversity of these cell lines. One is the A549 line, a cancer cell line representing type 2 alveolar cells (Foster et al, 1998); the other is the BEAS-2B cell line, an immortalized airway epithelial cell line (Fujisawa et al, 2000). Under unstressed conditions, all hFOXO proteins (hFOXO1, hFOXO3, hFOXO4, and hFOXO6) were mostly restricted to the cytoplasm in both cell lines (Figure 6A, G, H). All tested stressors induced nuclear translocation of hFOXO1 in A549 cells (Figure 6A-F). Quantitative analysis of nuclear translocation of hFOXO proteins revealed striking differences between the two cell lines (Figure 6G, H). hFOXO1 underwent nuclear translocation in response to all stressors in A549 cells (Figure 6G), but only in response to oxidative stress in BEAS-2B cells (Figure 6H). hFOXO3 underwent nuclear translocation in response to oxidative stress, starvation, and UV irradiation in A549 cells (Figure 6G) but not in response to any stressor in BEAS-2B cells (Figure 6H). hFOXO4 (Figure 6G, H) exhibited highly dynamic nuclear translocation following stress application in both cell lines, suggesting that different cell types of the airways respond differently to different stressors in terms of nuclear translocation of hFOXO proteins. For hFOXO6, the nucleus-to-cytoplasm ratio was already comparatively high under control conditions and changed only marginally upon stress application; given the very low expression of hFOXO6 in airway epithelial cells, these minor changes should be interpreted with caution and may in part reflect limited antibody specificity (Fig. 6G, H).

**Figure 6:**
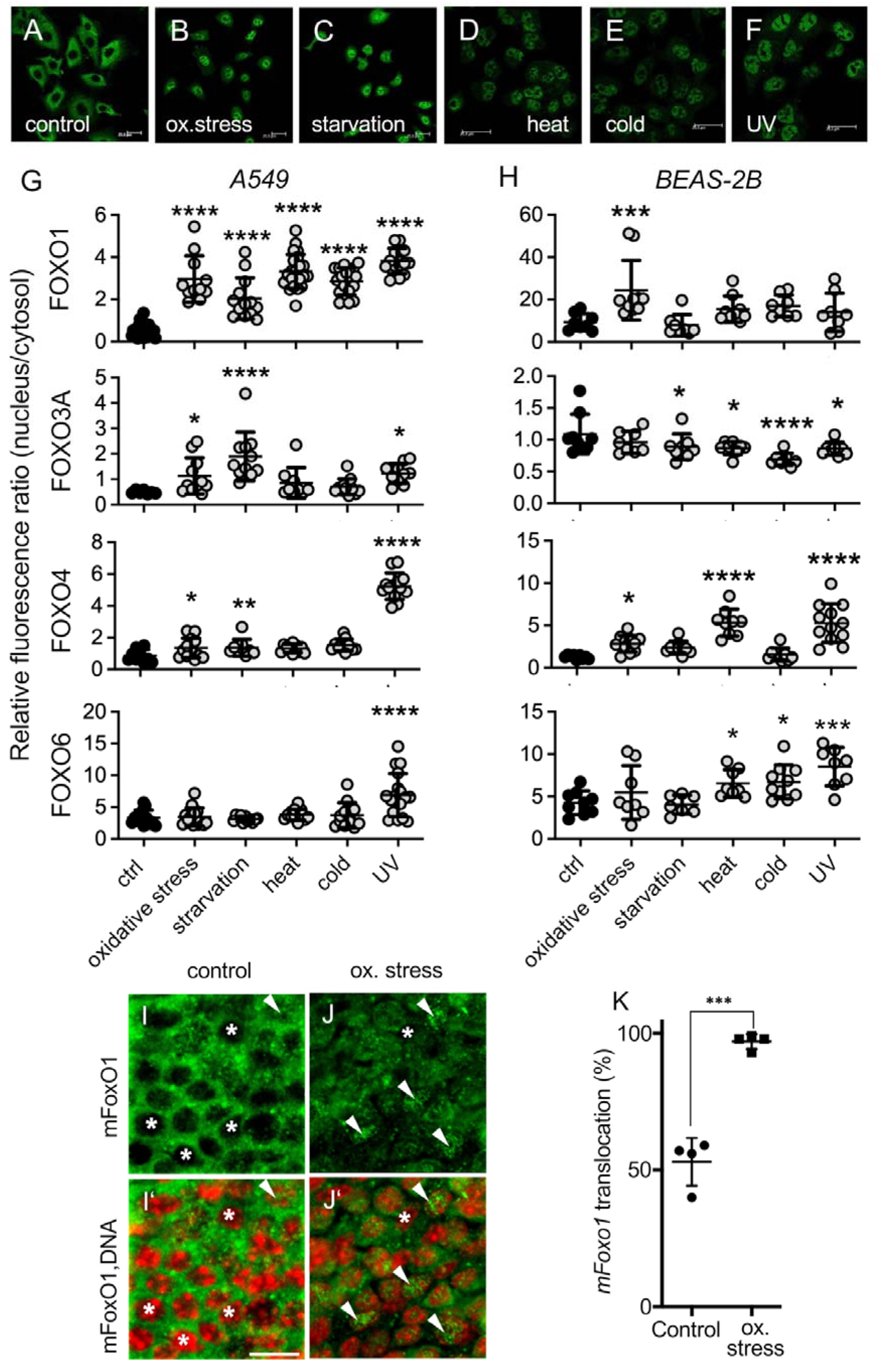
Stress-induced nuclear translocation of FoxO factors in AECs is conserved between species. A-F) Cultured A549 cells (human alveolar type II cells) were exposed to A) medium only (control) or different stressors, namely, B) oxidative stress induced by paraquat, C) starvation, D) UV light, E) cold stress, and F) heat stress. hFOXO1 was detected immunohistochemically. Scale bars are 25 µm. G-K) Nuclear translocation was quantified by determining the distribution of the antibody-linked fluorescence signal in the nucleus and cytoplasm of G) A549 and H) BEAS-2B cells. Cultured A549 and BEAS-2B (human bronchial epithelial cells) cells were exposed to medium only (ctrl), oxidative stress, starvation, heat stress, cold stress, and UV light. G, H) I mmunofluorescence staining revealed the cellular distribution of hFOXO1, hFOXO3, hFOXO4, and hFOXO6. The relative ratios of fluorescence intensities in the nucleus to those in the cytoplasm are shown. A ratio of 1 indicates that the fluorescence intensities in the cytoplasm and the nucleus are identical. The higher the ratio, the higher the fluorescence intensity in the nucleus. Each dot represents the mean value for a single biological replicate. I, J) Semi-intact and freshly explanted mouse tracheae were exposed to medium (control) or paraquat (oxidative stress). Nuclei were stained with Hoechst 33258 (red) and mFoxo1 was immunohistochemically stained (green). I ’, J’) The bottom images are merged. The scale bar is 10 µm. Asterisks indicate nuclear regions without mFoxo1 signal. Arrowheads indicate nuclear regions containing mFoxo1 signals. K) Quantification of nuclear translocation of mFoxo1 in murine tracheae exposed to paraquat. Points represent values obtained in independent experiments/individual animals (n=4). Significant differences compared with the respective control were calculated by one-way ANOVA with correction for multiple comparisons. *p<0.05, **p<0.01, ***p<0.001, and ****p<0.0001.

To validate this stress-induced nuclear translocation in an *in vivo* organ context, we challenged intact mouse tracheae with paraquat to induce oxidative stress. We focused on mFoxO1, which has shown a robust response to various stressors in cell culture experiments. Most of the mFoxO1 in untreated tracheae was in the cytoplasm and showed nuclear translocation in 52±4 % of the cells (Fig. 6I-K). After exposure to paraquat, a massive nuclear translocation of mFoxO1 (Fig. 6I-K) was observed in almost all cells (96±0.5 %; p<0.0001).

### 2.7 FoxO factors are reduced in airway epithelia of murine asthma models and in the sputum of asthmatic patients

So far we have shown that diverse environmental stressors consistently induce nuclear translocation of FOXO in the airway epithelia of flies, mice and humans, supporting a conserved cellular stress response. Consequently, we asked whether the lung disease that is characterized by intense stress sensitivity, namely asthma, shows altered FoxO signaling in the airways. We analyzed different mouse models of asthma (acute and chronic, Figure 7). In the acute asthma model, mice were sensitized to ovalbumin (OVA) (Figure 7A–D). We considered the distal and proximal airways separately by physically separating and isolating these airways from each other. There were significant differences between the proximal and distal airways. A distal airway was defined as the segment of a terminal bronchiolus that, starting at the bronchioalveolar duct transition, extended up to five alveoli along the proximal direction (Wegmann et al, 2005). In untreated control mice, the relative expression of mFoxo factors was higher in proximal airways than in distal airways, and these differences were statistically significant for mFoxo4 and mFoxo6 (Figure 7C, D). The transcript levels of mFoxo genes were always reduced in OVA-treated mice. The transcript levels of mFoxo1 (Fig. 7A) in the proximal airways, mFoxo4 (Figure 7C), and mFoxo6 (Figure 7D) in the proximal and distal airways were significantly lower in OVA-treated mice than in control mice. Finally, we compared whole lung transcript levels between control mice and acute and chronic asthma models (Figures 7E-H). mFoxo expression was strongly reduced in entire lung tissue from mice with chronic experimental asthma (Figures 7E-H). Relative to the transcript levels in lung tissue from healthy mice (set to 100 % in 7E-H), the expression level of mFoxo1 (Figure 7E), mFoxo3 (Figure 7F), mFoxo4 (Figure 7G), and mFoxo6 (Figure 7H) in lungs from mice with chronic experimental asthma went down to 50 %, 47 %, 40 %, and 30 %, respectively, and all these reductions were statistically significant.

**Figure 7:**
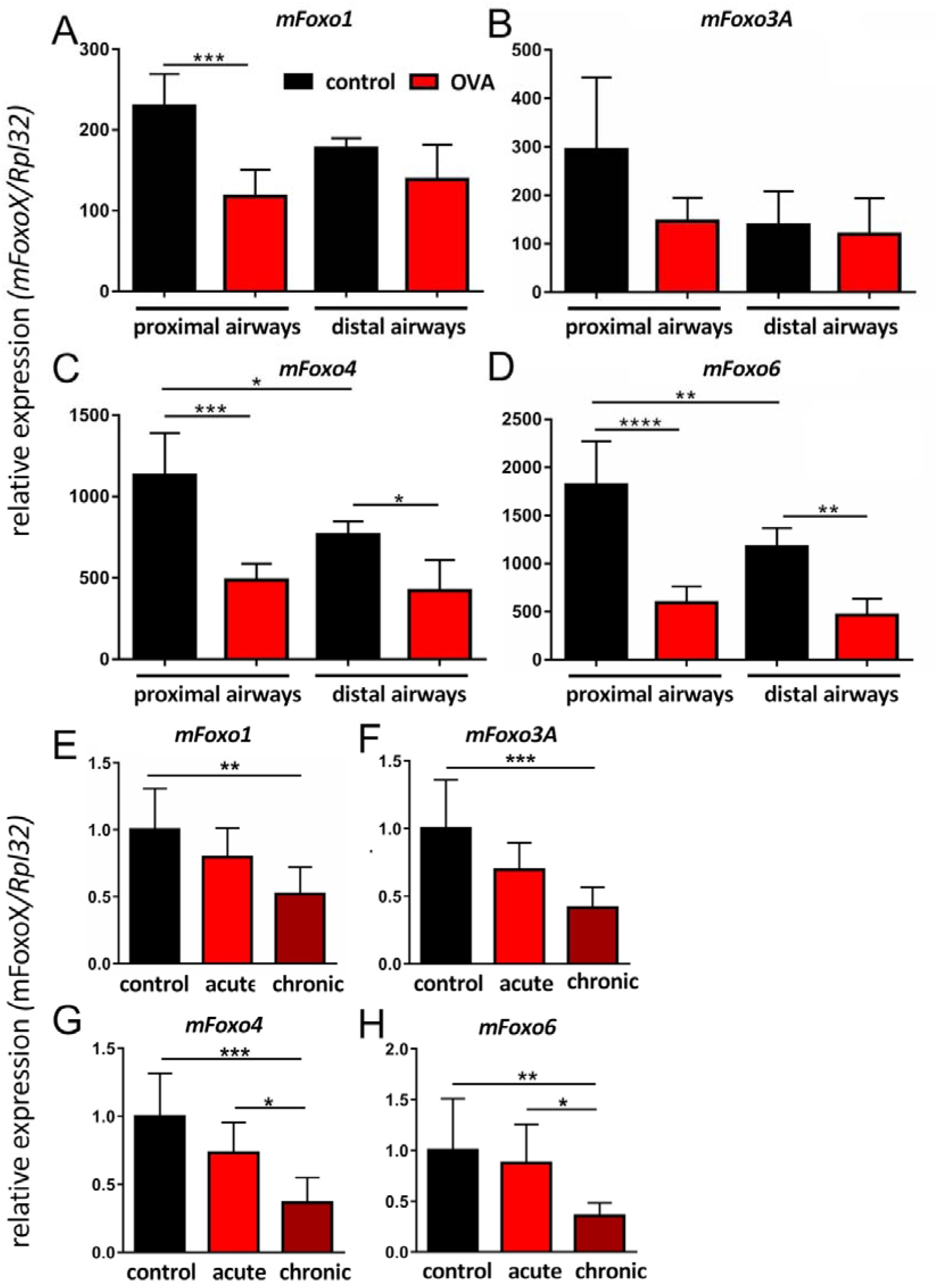
Expression of mFoxo1, mFoxo3, mFoxo4, and mFoxo6 in lung tissues of PBS-treated control mice and OVA-treated mice used as models of acute and chronic asthma. A-D) Micro-dissected airways of PBS-treated control mice and OVA-treated mice (acute asthma model) were separated into distal and proximal parts. A) mFoxo1, B) mFoxo3, C) mFoxo4, D) and mFoxo6 were expressed in both lung regions in both types of mice. In PBS-treated mice, expression was higher in proximal airways than in distal airways. Furthermore, expression in both distal and proximal airways was lower in OVA-treated mice than in control mice. However, this reduction was more apparent in proximal airways than in distal airways. In distal airways, expression of A) mFoxo1, B) and mFoxo3 did not significantly differ between control and OVA-treated mice. Data are presented as the mean and SD (n=5). Statistical analysis was performed using a one-way ANOVA. (E-H) Expression of the four mFoxo genes was monitored in lung tissues of PBS-treated control mice and OVA-treated mice used as models of acute and chronic asthma. Expression of E) mFoxo1, F) mFoxo3, G) mFoxo4, H) and mFoxo6 was reduced in lung tissue of the acute asthma mouse model and was decreased to an even greater extent in lung tissue of the chronic asthma mouse model. Data are presented as the mean and SD (n=4 mice per group). Statistical analyses were performed using a one-way ANOVA. *p<0.05, **p<0.01, ***p<0.001, and ****p<0.0001.

Finally, we wanted to see if these findings on the importance of FoxO factors in mouse and *Drosophila* models could also be observed in asthma patients. To this end, we compared the levels of hFOXO transcripts in sputum samples from asthmatic patients and healthy individuals. The levels of all four hFOXO factors were lower in sputum samples from asthmatic patients than in sputum samples from healthy controls (Figure 8A), and these differences were statistically significant for hFOXO1 and hFOXO3. We wanted to know whether hFOXO levels are also reduced in a matching well-defined experimental model of asthma. For this purpose, we used primary human epithelial cells from healthy donors and quantified the levels of hFOXO transcripts under control conditions and following Th2 priming by IL-4 treatment (Figure 8B). The transcript levels of all four hFOXO factors were strongly reduced in primary NHBEs stimulated with IL-4, and these differences were statistically significant for hFOXO1, hFOXO3, and hFOXO6, which showed reductions of 50 % or more (Figure 8B).

**Figure 8:**
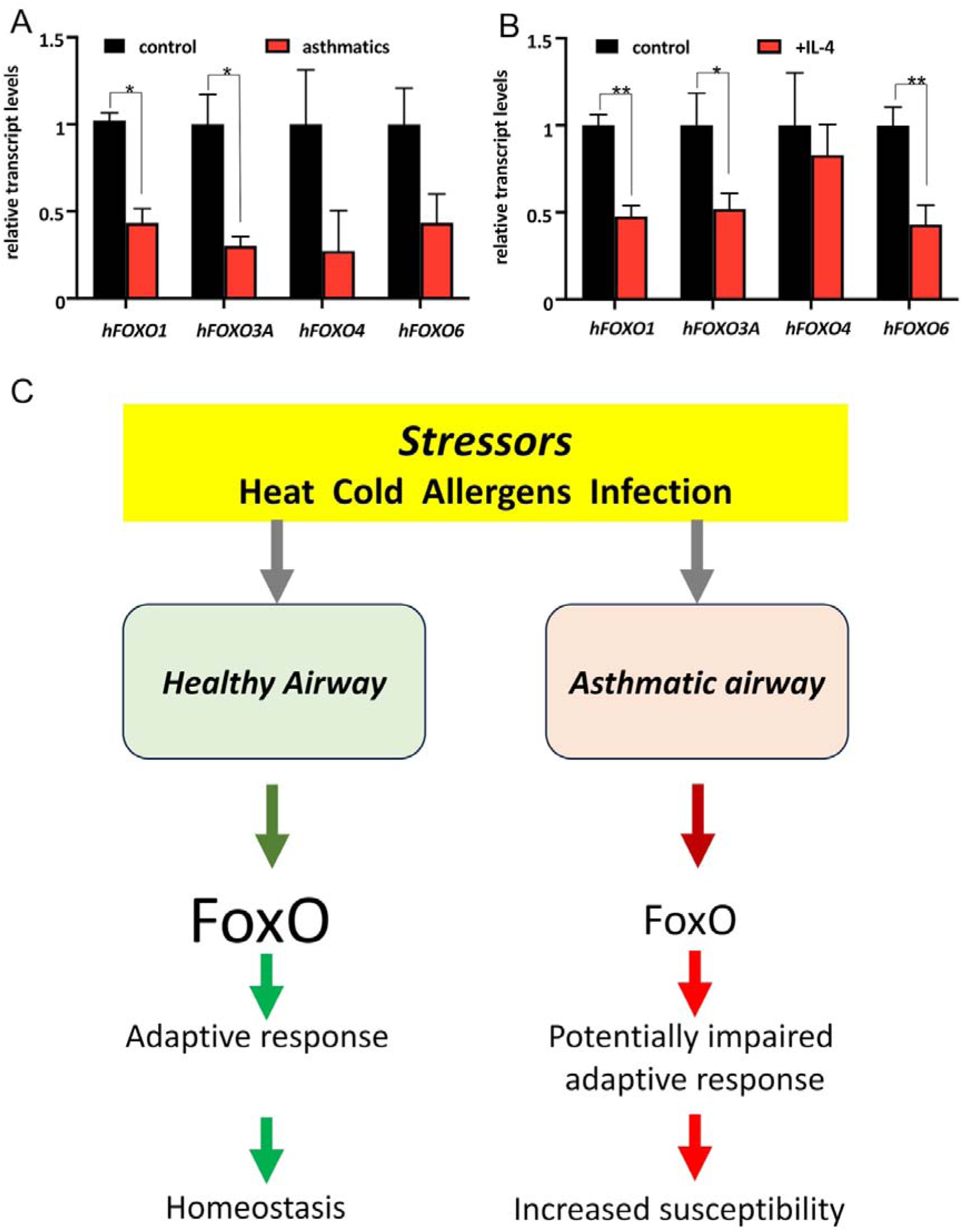
Transcript levels of hFOXO factors in sputum samples of asthmatic patients and primary AECs after Th1/Th2 priming. A) Relative levels of *hFOXO* factors in sputum samples of asthmatic patients and non-asthmatic controls were quantified by mRNA microarray analyses (n≥3). Statistical analysis was performed by the Mann-Whitney U test. B) NHBEs of healthy individuals were cultured under control conditions or treated with IL-4 to induce Th1/Th2 priming. Relative *hFOXO* transcripts were quantified by mRNA microarray analyses (n=6). Statistical analysis was performed by the Mann-Whitney U test. *p<0.05 and **p<0.01. C) Scheme summarizing the function of FoxO factors in a physiological state (left) or in a disease-associated non-physiological state (right).

We propose that appropriately regulated FoxO signaling may contribute to adaptive responses to a broad range of physiological stressors and thereby support airway homeostasis (Figure 8C). In asthmatic conditions, altered FOXO regulation may be associated with impaired stress adaptation, potentially contributing to exaggerated or maladaptive responses even under comparatively mild stress conditions (Figure 8C).

## 3 DISCUSSION

### 3.1 FoxO as a conserved epithelial stress sensor and its association with lung disease

The present work identifies a phylogenetically highly conserved mechanism of cellular response to stressors in airway epithelial cells (AEC). These epithelial cells orchestrate the response of the whole organ to various stressors and thus enable the maintenance of the functional integrity of the entire organ. We showed that this critical task of maintaining a homeostatic situation depends on the activity of FoxO factors in these epithelial cells. Consequently, deregulation of the FoxO system in AECs can be associated with pathological conditions. We have demonstrated reduced levels of FoxO factors in patients with asthma and in experimental murine asthma models. This association is particularly true for FoxO1, which appears to be affected in the inflamed lungs of both species and has been linked to various diseases (Peng et al, 2020). In addition, we have shown that loss of FoxO signalling drastically reduces resistance to airborne stressors. To demonstrate the latter, we deliberately used the simple *Drosophila* model, in which only one FoxO factor is present, and it also fulfills crucial functions in the respiratory epithelium (Wagner et al., 2021). The reduced hFOXO activity that we have shown to correlate with asthma in various models also characterizes other chronic lung diseases. In pulmonary hypertension, pathology correlates with decreased hFOXO1 expression (Savai et al., 2014). I t is also worth noting that deregulation of hFOXO1 in cells of the immune system, such as macrophages, may contribute to the pathogenesis of asthma (Chung et al, 2019). Reduced expression and activity of other hFOXO factors are also associated with lung diseases. For example, reduced hFOXO3 levels correlate with the development of COPD (Ganesan et al, 2013; Hwang et al., 2011; Pace et al, 2016) and IPF (Al-Tamari et al., 2018). This disease association is supported by the fact that hFOXO3 shows nuclear translocation in AECs from patients suffering from these diseases (Seiler et al., 2013). I t is also worth mentioning that loss of hFOXO1 and hFOXO3 function is associated with several forms of lung cancer (Gao et al, 2018; Mikse et al., 2010; Wu et al, 2022). Although our findings support the concept that FOXO responsiveness to epithelial stress is evolutionarily conserved, our data do not imply that identical FOXO-dependent transcriptional programs operate across species. Rather, we propose that conservation primarily resides at the level of stress sensing and adaptive signal integration, whereas the downstream transcriptional implementation is likely shaped by the distinct physiological requirements and evolutionary history of insect and mammalian airway systems

Collectively, our findings identify FoxO as a central regulator of epithelial stress adaptation that enables the maintenance of airway homeostasis under chronic environmental stress.

### 3.2 Reduced FoxO activity, stress sensitivity and impaired epithelial repair

Using the *Drosophila* system, we showed that reduced dfoxo levels and disease-related increased sensitivity to various stressors are causally linked. Following this line of thought, the reduced ability to respond appropriately to multiple stressors due to reduced hFOXO expression, and presumably reduced FOXO activity, could lead to disease development in patients. Studies showing that FoxO factors orchestrate epithelial homeostasis, particularly in response to respiratory stressors and infections, are consistent with this interpretation (Gimenes-Junior et al., 2019; Pomies et al, 2016). Furthermore, skin epithelia defective in hFOXO1 expression have reduced repair capacity (Mori et al, 2014), a finding that may also apply to the lung epithelium. Further evidence supports the interpretation that hFOXO factors play a central role in various lung diseases. The main observation is that a reduced ability to repair damaged epithelia can be a decisive driver of a wide range of lung diseases. This interpretation applies to chronic lung diseases such as COPD and IPF (Barnes et al, 2021), in which this reduced ability to repair can be triggered by the disease or acquired during the disease. The significance of hFOXO factors for the genesis of these diseases is an exciting topic for future research. Nevertheless, many lines of evidence already suggest that hFOXO factors are potential targets for specific interventions.

### 3.3 FOXO signalling in type-2 inflammatory airway environments

Our studies on the Th2 polarisation of NHBEs (Zissler et al, 2016) indicate a particularly close relationship between asthma and hFOXO factors. The polarization of primary epithelial cells induced by IL-4 treatment reduced the expression of all hFOXO factors, as seen in asthma patients. This observation suggests that Th2-associated epithelial remodeling is accompanied by reduced FOXO expression. Together with the findings in asthma patients, this supports a link between type-2 inflammatory airway environments and altered FOXO signaling. Whether reduced FOXO activity directly contributes to impaired epithelial repair or represents a consequence of chronic inflammation remains to be determined.

### 3.4 dFoxo-dependent transcriptional programs in the airway epithelium

To gain mechanistic insight into the role of dFoxo in airway epithelial homeostasis, we performed transcriptomic analyses of isolated *Drosophila* airways following dfoxo loss- and gain-of-function. These analyses revealed a broad reciprocal regulation of genes involved in epithelial maintenance, stress protection, metabolic adaptation, and developmental processes. In particular, dfoxo deficiency reduced the expression of genes associated with glutathione-dependent detoxification, xenobiotic metabolism, amino acid metabolism, and multiple stress-response pathways, while simultaneously increasing the expression of genes involved in DNA replication, cell-cycle progression, and developmental signaling. Conversely, airway-specific dfoxo overexpression promoted the expression of genes associated with cuticle organization and tissue maintenance. Together, these findings suggest that dFoxo functions as a central regulator of airway epithelial resilience by promoting maintenance and stress-protective programs while restricting excessive growth-associated activity.

Among the dFoxo-regulated genes, heat shock proteins were of particular interest. Heat shock proteins are essential components of cellular stress protection and have previously been implicated in asthma and other chronic lung diseases (Donovan & Marr, 2016; Hou et al, 2011; Son et al, 2018). The reduced expression of several HSPs in dfoxo-deficient airways is therefore consistent with the increased stress sensitivity observed in these animals and supports the notion that HSPs represent one important downstream component of the broader dFoxo-dependent stress-response network (Donovan & Marr, 2016; Kim & Koh, 2017; Tower, 2011).

### 3.5 Stressor-, FOXO- and cell type-specific responses across model systems

The potential disease relevance of deregulated FOXO expression in AECs is a direct consequence of the central physiological role of these transcription factors in adaptation to changing conditions, whether externally or internally induced. Because FoxO factors in airway epithelial cells respond to various stressors, they are ideal signaling hubs for adapting cellular stress responses. The phylogenetic conservation of this stress sensor system enables analysis of its general properties in the simple *Drosophila* model. Here, we could also determine the causal relationship to reduced stress resistance.

In this study, we have selected a significant number of stressors that, at first glance, do not appear to be equally suitable across all systems but can provide us with information about the general stress-response behavior of the AEC. For example, we chose paraquat as an oxidative stress generator because it can be used equally well in all systems ( *Drosophila*, mouse, cell culture). Although heat and cold are extremely relevant stressors in *Drosophila*, they are only of limited significance in organisms of the same temperature, such as mice and humans. However, this does not apply to respiratory air, which can introduce such temperature differences into the AEC. In this context, we must realize that not all stressors are mediated via the same signaling pathway to FoxO activation. There is a differential response to stressors, particularly among the various hFOXO factors in humans. In contrast to the simple system of a single FoxO factor in *Drosophila*, the human system is complicated because four FoxO factors play this role. The presence of multiple FoxO factors means that the vertebrate system is characterized by high complexity, redundancy, and overlapping functions. In AECs, the four hFOXO factors exhibited distinct response spectra to different stressors. HFOXO1 responds strongly to various stressors with nuclear translocation. Our hypoxia experiments further support the concept of FOXO factors as conserved airway stress sensors. Like the activation of dfoxo observed in *Drosophila* airways, hypoxia induced a pronounced nuclear translocation of hFOXO1 in human airway epithelial cells. This finding suggests that oxygen deprivation represents an evolutionarily conserved trigger of FOXO activation in respiratory epithelia. Interestingly, hypoxia also induced the expression of the pro-inflammatory mediators CCL20 and CXCL8, indicating that FOXO activation occurs in parallel with epithelial stress and inflammatory responses. Whether FOXO signaling directly contributes to the regulation of these cytokines or primarily serves to protect epithelial integrity during hypoxic stress remains to be determined. Nevertheless, these observations further support the notion that FOXO factors function as central integrators of environmental stress signals in airway epithelial cells. For the hFOXOs hFOXO3 and hFOXO4, the picture is somewhat more differentiated: we observed responses to fewer stressors in each case, and these responses also vary strongly between cell types. hFOXO6, on the other hand, generally shows low activation levels but is also expressed at low levels and shows strong nuclear localization even under control conditions, making staining prone to background artifacts. Despite the wealth of information and structural differences leading to different activation modes, our knowledge of the interplay of hFOXO factors still needs to be more comprehensive (Potente et al, 2005; Spreitzer et al, 2022). This lack of information concerns different targets and aspects, such as the complementarity of the action spectra or functional substitution through compensatory effects. Together, these findings indicate that FOXO-dependent stress responses are both cell type- and FOXO-specific in the human airway epithelium. Although FOXO1 has been described as a FOXO3-responsive gene in certain cellular contexts, the relationship between FOXO3 nuclear localization and FOXO1 regulation in airway epithelial cells remains unclear and may depend on additional context-specific regulatory mechanisms.

### 3.6 Conclusions

Our results show that FoxO factors in the airway epithelium respond to numerous stressors by translocating to the nucleus. In mammals, this response is FoxO factor- and cell type-specific, although there is considerable overlap. This observation suggests that the mammalian lung has a highly complex, region-specific stress-sensing system based on FoxO factors. Furthermore, we have shown that a lack of dfoxo expression substantially enhances stress sensitivity, which may underlie the development of several chronic lung diseases. Taken together, our findings support a model in which FOXO factors act as central regulators of airway epithelial stress adaptation. Rather than functioning solely as immune regulators, FOXO proteins appear to contribute to the resilience of airway epithelia against diverse environmental challenges. Impaired FOXO signaling may therefore reduce the capacity of airway tissues to maintain homeostasis under chronic stress conditions and could represent an important component of disease susceptibility in chronic inflammatory lung disorders.

### 3.7. Limitations of the study

This study aimed to identify phylogenetically conserved mechanisms of FOXO activation in airway epithelial cells by integrating complementary experimental systems in *Drosophila*, mouse, and human samples. Although this comparative approach offers a comprehensive perspective on conserved FOXO-dependent stress responses, each model system addresses distinct aspects of airway biology and possesses inherent limitations. As a result, certain mechanistic questions could not be fully resolved within a single experimental system. Specifically, the human studies were limited by the availability of patient material and therefore establish associations, rather than causal relationships, between altered FOXO signaling and airway disease. Furthermore, it is important to acknowledge that elucidating the molecular nature of FoxO modification, for instance by Western blotting or proteomic approaches, as well as defining the contribution of insulin signaling to the phenotypes described here, lies beyond the scope of the present study and will require dedicated future investigations. This also applies to cell-type-specific genetic studies and in-depth comparative studies of evolutionary conservation of the observed results.

While these findings establish an evolutionarily conserved role for FOXO signaling in airway epithelial stress adaptation, further research is necessary to define its causal contribution to chronic airway disease. Employing tissue-specific genetic perturbation in physiologically relevant mammalian airway models will be essential to determine whether reduced FOXO activity directly impairs epithelial stress adaptation or is a consequence of chronic inflammation. Such investigations will further elucidate the role of FOXO-dependent pathways in disease susceptibility, progression, and tissue repair. Building on these points, we identify three key areas for future work. First, conditional, cell type-specific deletion of individual Foxo factors (particularly Foxo1 and Foxo3) in the murine airway epithelium, combined with controlled exposure to the stressors used here, would determine whether reduced FoxO activity is a cause or a consequence of impaired epithelial stress adaptation. Second, primary air-liquid interface cultures from patients and matched controls would enable assessment of subcellular localization of the hFOXO factors in a human system that maintains epithelial differentiation (something not possible in sputum). Thirdly, FoxO activity should be monitored by target gene signatures or reporter systems rather than by expression levels per se, as our data clearly show that the two are not necessarily the same. Finally, molecular characterization of stressor-specific FoxO modifications would link the conserved activation mechanism described here to its functional outcomes in chronic airway disease.

## 4 MATERIAL AND METHODS

### 4.1 *Drosophila* stocks, maintenance, and stress assays

Stocks and transgenic crossings were raised on standard cornmeal-agar medium at 21 °C at a relative humidity of at least 60 % in a 12 h:12 h light/dark cycle as described earlier (von Frieling *et al*, 2020; Wagner *et al*., 2021). The following strains used were supplied by the Bloomington and Kyoto/DGGR Stock Center or by scientists of the *Drosophila* research community: *CantonS* (BL1), *w^1118^* (BL3605), *Drs-GFP* (BL55707), UAS-*bsk^DN^* (BL6409), UAS-gRNA-dfoxo (BL82742), UAS-Cas9.P (BL54592), *btl*-Gal4 (Christian Klämbt, Münster, Germany), *ppk4*-Gal4 (Mike Welsh, Iowa, USA), UAS-*foxo-gfp* (Wagner *et al*., 2009), UAS-*foxo* (BL9575), UAS-*foxo*-TM (Marc Tatar, Brown University, USA), *foxo^21/21^*(Ernst Hafen, Zuerich, Switzerland) (Junger *et al*., 2003), UAS-*relish-yfp, LDH-Gal4/UAS-nGFP* (Pablo Wappner, Buenos Aires, Argentina, (Lavista-Llanos *et al*., 2002)), and UAS-*dorsal-gfp* (Tony Ip, Massachusetts, USA). To induce expression of either genes of interest or GFP fusion proteins in larval airways, we used the binary Gal4/UAS expression system (Brand & Perrimon, 1993). *dFoxo* overexpression experiments were performed using the Target system employing the temperature-sensitive tubP-Gal80ts element (McGuire *et al*, 2004). Therefore, controls were maintained at 18°C, whereas activation was achieved by shifting the temperature to 30°C for 24 h.

For all stress tests, early 3^rd^-instar larvae were used. Starvation, irradiation, and oxidative stress experiments were done in PBS at 21°C. Starvation experiments proceeded by incubation on PBS-soaked filters for 12 h. For irradiation experiments, larvae were exposed to UV light (254 nm, distance 1 cm) for 15 min, followed by incubation in a medium for up to 2 h. Oxidative stress was induced using the chemical paraquat (Methyl viologen dichloride hydrate, Sigma-Aldrich, Steinheim, Germany) at a 100 mM working concentration in an enriched medium solution. Larvae were incubated in the paraquat solution for 1 h and then transferred to medium for an additional hour. Temperature experiments were performed within the medium. Animals subjected to heat shock were incubated at 37°C for 30 min; translocation experiments were carried out immediately, or the larvae remained for 90 min at room temperature. Cold treatment was performed at 4°C for 2 h. Hypoxia (1% O_2_) was applied for 4 h under otherwise normal conditions.

Hypoxia-induced larval escape assays were performed as shown (Kallsen et al., 2015). For the hypoxia recovery assay, animals were briefly (15 s) anesthetized with N_2,_ and the time to recovery was measured. For the desiccation assay, animals were placed into empty vials under constant conditions, and survival was monitored. For chronic cigarette smoke exposure, animals were subjected to daily doses of cigarette smoke, and their survival was also monitored daily. For hypoxia experiments, animals were subjected to 5% O_2_ for the indicated times.

Microscopic analyses in *Drosophila* were performed in vivo using early 3^rd^ instar larvae with an SZX16 stereo microscope and a DP72 camera (Olympus, Hamburg, Germany), and a Zeiss 880 CLSM. GFP signals were detected with an Ex 460-490nm.

### 4.2 *Drosophila* transcriptomics

Tracheae were manually isolated from 3^rd^ instar larvae and processed as described earlier (Bossen *et al*, 2019; Kallsen *et al*., 2015; Prange *et al*., 2018). The dissected tissue was suspended in RNA Magic (Bio Budget) and homogenized in a bead mill using 1.4 mm zirconia beads (Biolabproducts, Bebensee, Germany). Total RNA input quality was evaluated on a TapeStation 4200 (Agilent, USA), and all samples showed a RIN score > 8. Samples were quantified with a fluorometric dye (Quant-IT, Thermofisher, USA), and between 187 and 500 ng per sample were used as input for the TruSeq stranded mRNA library kit (I llumina, USA) following the manufacturer’s manual. Resulting libraries showed a fragment size distribution of around 350 bp and were sequenced on a HiSeq 4000 (I llumina, USA) with 50bp single end reads. Differential expression of sequencing data (GSE176540) was performed with the CLC genomic workbench software (http://www. clcbio.com/products/clc-genomics-workbench) (Qiagen, Hilden, Germany), using the *Drosophila* melanogaster reference genome (Release 6) (Hoskins et al, 2015). GO analyses were conducted using the g:Profiler tool (Raudvere et al, 2019).

### 4.3 Cell culture (human-derived cell lines) and stress assays

The alveolar epithelial cell line A549 and the bronchial epithelial cell line BEAS-2B were cultured in RPMI (Biochrom, Berlin, Germany) supplemented with 10% fetal calf serum, 100 U/ml penicillin, and 100 µg/ml streptomycin at 37°C with 5% CO_2_. Stress experiments were performed in µ-Slides 8-well ibiTreat (Ibidi, Martinsried, Germany) with 2,5 x 10^4^ cells per well; cells adhered to the slides overnight. Medium was replaced with either pure RPMI (unsupplemented control), RPMI with paraquat [20 mM] (oxidative stress), or PBS (starvation) and incubated for 1 h at 37°C with 5 % CO_2_. Cold and heat stress were induced for 1 h each at 41°C and 4°C, respectively. Exposure to UV light (254 nm, distance 1 cm) took place for 15 min, followed by incubation up to 1 h. Hypoxic conditions were established in a supplemented medium for 2 h at 5% O_2_ by discharging N_2_ into a gas-tight chamber. Immediately afterward cells were fixed with 3 % paraformaldehyde, permeabilized with 0,25 % Triton-X/PBS, blocked with 10 % BSA/PBS and incubated with primary antibodies: anti-FOXO1 antibody (rabbit polyclonal, ChIP grade), anti-FOXO3 antibody (rabbit polyclonal), anti-FOXO4 antibody (rabbit polyclonal); (abcam, Cambridge, UK) and anti-FOXO6 antibody (rabbit polyclonal; Biorbyt, Cambridge) or anti-FOXO3 antibody (rabbit polyclonal); (Thermo Sci. / Pierce Biotech, Rockford, USA), secondary antibody: Alexa Fluor 488 goat anti-rabbit IgG (Invitrogen, Karlsruhe, Germany). Nuclei were stained with Bisbenzimide H 33342 (Sigma-Aldrich, Deisenhofen, Germany).

Human cells (A549 and BEAS-2B) were investigated using the TCS Sp5 Inverted Confocal laser scanning microscope (Leica Microsystems, Wetzlar, Germany) and a 20x /0.70 HC Plan Apochromat, oil-immersion objective. Antibody and Hoechst signals were detected with excitation lines of 488 and 405 nm, respectively. All editing procedures were carried out with the Leica LAS AF Lite program. Quantification of the nuclear translocation in A549 and BEAS-2B was performed using ImageJ software (Wayne Rasband, NIH). Randomly selected cells were photographed, and all images were taken with constant settings. Considering the area size, the average fluorescence intensity of the nucleus and cytoplasm was determined. The nucleus-to-cytoplasm ratio is shown graphically and compared with that of untreated control cells.

Relative quantification of human cytokine transcripts of CCL20 and CXCL8 (IL8) was performed by real-time PCR. Total RNA (A549 cells; standard incubation control and after 1, 2, 4 h under hypoxic conditions) was extracted using the NucleoSpin RNA Kit (Macherey-Nagel, Düren, Germany) according to the manufacturer’s instructions. 500 ng of each RNA sample was transcribed into cDNA, using Oligo dt 12-18 primer mix, RNAse OUT, dNTP mix, and SuperScript reverse transcriptase (Invitrogen, Darmstadt, Germany)). The human HPRT gen served as reference für relative expression. Statistical analysis: ratio one-way Anova, CCL20: n=7, CXCL8 n=6.

### 4.4 Detection of mFoxO1 nuclear translocation in AECs of the murine trachea

Three-month-old C57BL/6 female mice were used. All experiments were performed in accordance with the government guidelines for animal welfare of the Land Schleswig-Holstein. Mice were killed by an overdose of isoflurane, and the trachea was explanted, opened along the tracheal muscle, and cut in half. One half was incubated for 30 min at 37 °C in 100 mM paraquat (Sigma-Aldrich, Taufkirchen, Germany) prepared in HEPES-buffered Ringer solution. The other half was incubated in HEPES-buffered Ringer solution lacking paraquat. Thereafter, the tissue was fixed with PBS containing 4 % paraformaldehyde and processed for immunohistochemistry with an anti-FoxO1 antibody (1:1000 in TBS, Thermo Fisher Scientific, Rockford, USA) at room temperature overnight followed by Alexa Fluor 555-conjugated donkey anti-rabbit IgG (1:500 in TBS, Invitrogen, Karlsruhe, Germany) for 1 h at room temperature and then Hoechst 33258 (1:10,000 in TBS; Sigma-Aldrich GmbH, Taufkirchen, Germany) for 10 min. Three-dimensional confocal image stacks of the epithelial layer were acquired and processed using the ImageJ “Cell Counter” Plugin. AECs were counted in 4 stimulated and 4 control pieces of the trachea. Nuclear regions were identified using DAPI signals, and the regions of the nucleus versus the regions of the cytoplasm were measured. Approximately 200 nuclei were analyzed per piece.

### 4.5 Quantification of gene expression in mouse lung tissue

Animal experiments were performed using female C57BL/6J (Charles River, Sulzfeld, Germany) mice for expression analysis in proximal and distal airways and for expression analysis in acute and chronic asthma models. All animal studies were approved by the local animal ethics committee (for the acute asthma model: 13-3/20, for the chronic asthma model: 55-6/20; Behörde das Ministerium für Landwirtschaft, ländliche Räume, Europa und Verbraucherschutz Schleswig-Holstein).

Mice were subjected only to the standard mouse protocol for acute experimental allergic asthma (Bulek et al, 2009), while the other mice were exposed to the protocols for both acute and chronic experimental allergic asthma (Wegmann *et al*., 2005). I n brief, mice were first systemically sensitized against ovalbumin (OVA) by intraperitoneal (i.p.) injection of 10 µg dissolved OVA in 100 µl PBS and 100 µl aluminium hydroxide as adjuvant on day 1, 14 and 21. On days 26-28, animals were exposed to OVA aerosol (1% OVA dissolved in PBS) in an airtight chamber for 20 min. For the chronic asthma model, the sensitization was continued twice weekly for 12 weeks. Control animals received an i.p. injection with PBS on the same days and were treated with a PBS aerosol likewise. Lungs were filled with 600 µl RNAlater (Thermo Scientific, Darmstadt, Germany), dissected, and lungs were divided into proximal and distal parts if necessary. The tissue was transferred into RNA lysis buffer, snap-frozen in liquid nitrogen, and homogenized mechanically with the addition of liquid nitrogen. RNA isolation was performed from 30 mg using the RNeasy Micro Kit (Qiagen, Hilden, Germany) according to the manufacturer’s instructions. RNA concentration and purity were determined using a nanophotometer. Reverse transcription of the isolated RNA into cDNA was performed using the Maxima First Strand cDNA Synthesis Kit (Thermo Scientific, Darmstadt, Germany) according to the manufacturer’s instructions. Gene expression was determined by qRT-PCR on Light Cycler 480II (Roche, Mannheim, Germany) using 2.5 μl cDNA, SYBR Green Mastermix (Roche, Mannheim, Germany), and 10 pmol/μl of each primer. Expression of the gene rpl32 was used as a reference. The sequences of the primers were as follows: rpl32: for 5’-AAAATTAAGCGAAACTGGCG-3’, rev 5’-GAAGATGGTGATGGGCTTCC-3’; mfoxO1: for 5’-CTTCAAGGATAAGGGCGACA-3’, rev 5’-GACAGATTGTGGCGAATTGA-3’; mfoxO3a: for 5’-GCTAAGCAGGCCTCATCTCA-3’, rev 5’-TTGTGTCAGTTTGAGGGTCT-3’; mfoxO4: for 5’-TGTGCTCGCATCTCCTACTG-3’, rev 5’-GACTCAGGGATCTGGCTCAAA-3’; mfoxO6: for 5’-CAAGAGACTCACGCTCTCGC -3’, rev 5’-GCCGAATGGAGTTCTTCCAGC-3’.

### 4.6 Human study subjects, lung function testing, sputum sampling, and primary cell culture

Healthy subjects (n=12) and mild, well-controlled, allergic asthma patients along the GINA definition in season (n=9) in good health (FEV1% >70%) with a history of clinically significant hay fever during the grass-pollen season for more than two years were included. For asthmatic subjects, a GINA classification was performed. All subjects completed a Rhinoconjunctivitis Quality of Life Questionnaire (RQLQ), a lung function test, and a hypertonic saline sputum induction, performed in the Allergy Section, Department of Otolaryngology, TUM School of Medicine. Each participant provided written informed consent. This study was approved by the local ethics committee (5534/12).

Baseline lung function was evaluated using a calibrated handheld pulmonary function testing device (Jaeger SpiroPro; Würzburg, Germany). The following parameters were recorded: VC, FEV1, FEV1/VC, and maximum expiratory flow 25% (MEF 25%). Bronchodilatator reversibility was tested after 400 µg of salbutamol.

Participants first inhaled salbutamol and consecutively nebulized hypertonic saline at increasing concentrations of 3%, 4%, and 5% NaCl every 7 min according to previous publications. During this procedure, participants cleaned their noses and rinsed their mouth to reduce squamous epithelial cells in the samples. Sputum was processed within one hour of collection. The selected sputum plugs, which contained as little saliva as possible, were placed in a weighed Eppendorf tube and processed with 4x weight/volume of sputolysin working solution (Merck, Darmstadt, Germany). Afterward, 2x weight/volume of phosphate-buffered saline (PBS) was added. Samples were filtered through a 70-µm mesh and centrifuged for 10 min at 790 × g without brake to remove the cells. Appropriate sputum samples were obtained from 12 healthy controls and 9 asthma patients during the grass-pollen season.

Primary NHBEs (Lonza, Walkersville, MD) of six genetically independent donors were grown as monolayers in 100% humidity and 5% CO_2_ at 37 °C in serum-free defined growth media (BEGM, Lonza). NHBEs (passage 3) were used at about 80% confluence in 6-well plates. To avoid gene expression changes or influences on the IL-4 signaling induced by growth factors in the BEGM medium, cells were rested in basal medium (BEBM) for 12 h, then stimulated with recombinant human IL-4 at 50 ng/mL (R&D Systems, Minneapolis, MN) in BEBM medium for 6 h or BEBM medium as control condition. For RNA analysis, harvested cells were lysed in RLT buffer (Qiagen, Hilden, Germany) containing 1% beta-mercaptoethanol (Roth, Karlsruhe, Germany) directly in the cell culture well.

Total RNA was extracted using RNeasy Mini Kit (Qiagen, Hilden, Germany) with on-column DNase digestion (Qiagen) for avoiding DNA contaminations. RNA quantification was performed by ultraviolet–visible spectrophotometry (Nanodrop Technologies, Wilmington, DE), for assessment of the RNA integrity by the RNA 6000 Nano Chip Kit with the Agilent 2100 Bioanalyzer (Agilent Technologies, Waldbronn, Germany), and samples with RIN number above 7.5 were used for micro-array analysis. Total RNA was amplified and Cy3-labeled by using the one-color Low Input Quick Amp Labeling Kit (Agilent Technologies) according to the manufacturer’s protocol. Hybridization to SurePrint G3 Human Gene Expression 8x60K Microarrays (Agilent Technologies) was performed with the Gene Expression Hybridization Kit (Agilent Technologies) as described before.

Data import was performed using a standard baseline transformation to the median of all values, including log transformation and computation of fold changes. Subsequently, a principal component analysis (PCA) was conducted and revealed a homogeneous component distribution. Compromised array signals (array spot is non-uniform if pixel noise of feature exceeds threshold or above saturation threshold) were excluded from further analysis. For this analysis, we set the normalized values of each healthy donor in sputum samples to 1 and calculated the difference of FoxO1, FoxO3, FoxO4, and FoxO6 expression in sputum of the asthma patient. For analysis of FoxO1, FoxO3, FoxO4, and FoxO6 expression in primary normal human bronchial epithelial cells (NHBEs), we set the normalized values of medium conditions (in NHBEs) to 1 and calculated the difference in abundance of the IL-4 stimulated condition. The data discussed in this publication have been deposited in NCBI’s Gene Expression Omnibus (GEO) and are accessible through GEO Series accession number GSE167225.

For RNA quality assessment, RNA integrity numbers (RIN) were measured via Agilent 2100 Bioanalyzer (Agilent Technologies, Santa Clara, CA, USA). After quality assessment, isolated total RNA was subjected to reverse transcription using a high-capacity cDNA kit (Applied Biosystems, USA), following the manufactureŕs instructions. Real-time PCR profiles were visualized using the commercially available FastStart Universal SYBR Green Mastermix (Roche, SUI) and Qiagen human Primer Assays (Qiagen, Hilden, Germany) specific for *FoxO1* and *FoxO3* (the latter only in NHBEs) and quantified by the ViiA 7 Real-Time PCR System (Applied Biosystems, USA). The amount of *FoxO1* and *FoxO3* mRNA expression was normalized with endogenous control *TFRH*, *GAPDH*, and *ß-actin* (housekeeping gene index, DCt values) and the relative quantification and calculation of the range of confidence was performed using the comparative threshold cycle (2^-DDCt^) method (relative gene expression). All amplifications were carried out at least in duplicate.

### Statistical analysis and data availability

RNA-seq and microarray datasets have been deposited in GEO under accession numbers GSE176540 and GSE167225. All other data supporting the findings of this study are available from the corresponding author upon reasonable request.

Investigators were blinded during analysis of human samples. Blinding was not applied to *Drosophila* and mouse experiments because experimental groups were predefined and objective quantitative readouts were used. Non-parametric Mann–Whitney U tests were used to determine significant differences between patient groups. All statistical tests were performed two-sided. Unless otherwise stated, n refers to independent biological replicates. Technical replicates were averaged before statistical analysis. No statistical methods were used to predetermine sample size. Sample sizes were based on previous experience with comparable experiments and published studies. Usually, samples were processed in parallel without random allocation.

Survival data were plotted as Kaplan–Meier curves, and survival of experimental and control genotypes was compared by the log-rank (Mantel–Cox) test, with each fly treated as an individual observation. Usually, flies were used in batches of 10 individuals.

## AUTHOR CONTRIBUTIONS

T.R., H.H., and P.I .P. conceived the project. T.R., H.H., P.I.P., I.B., K.U., C.S.W., P.K. wrote the manuscript. K.U., J.B., X.N., U.Z. R.P., C.F., M.P., C.V., C.W., S.F., and A.A. performed all experiments, analyzed the data, and made the figures.

## ACKNOWLEDGMENTS

This work was supported by grants from the Deutsche Forschungsgemeinschaft (DFG) as part of the CRCs TR22 (TPA7) and 1182 (TPC2) and INST 257/591-1 FUGG, as well as from the University of Kiel in the program of the LeibnizCampus EvoLung.

## DATA AVAILABILITY STATEMENT

The data that support the findings of this study are available from the corresponding author upon reasonable request.

