## Supplementary figures and images for "FoxO factors preserve airway epithelial homeostasis by coordinating adaptive stress responses"

### Fig. 4 Supplemental Fig. 1

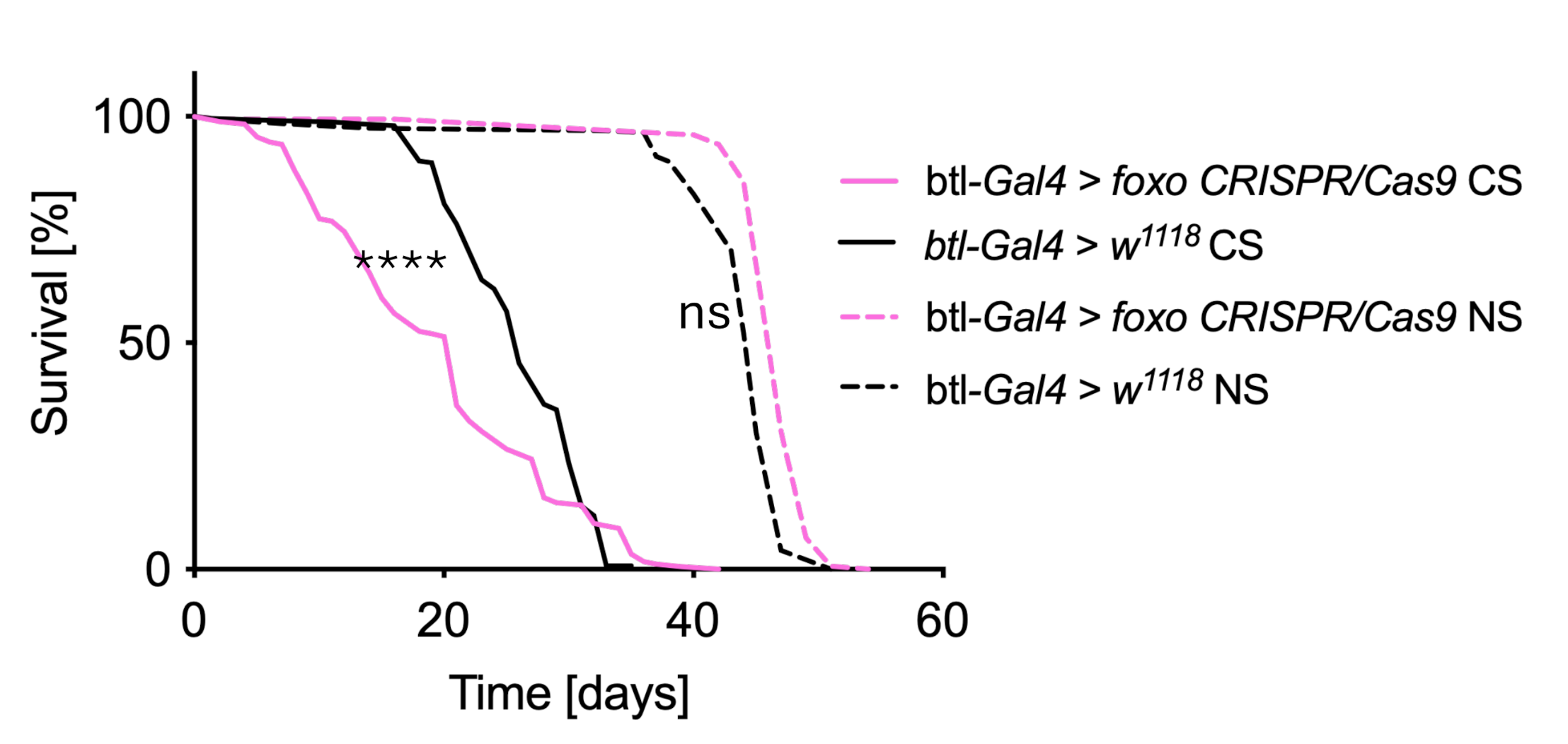
