## Supplementary material for "FoxO factors preserve airway epithelial homeostasis by coordinating adaptive stress responses": Fig.6 Supplemental Fig. 1

A549

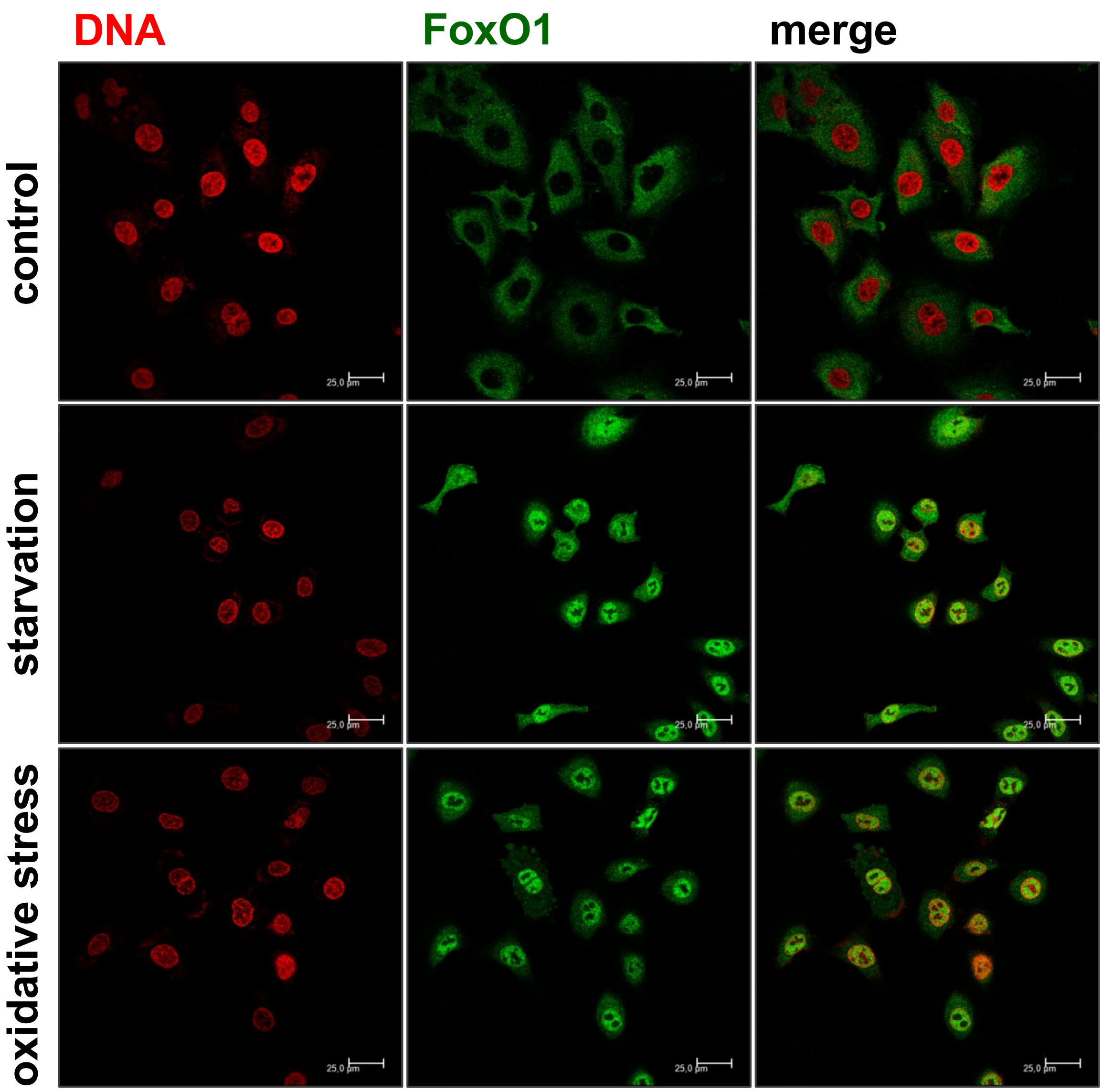
DNA
FoxO1
merge
control
starvation
oxidative stress
25,0 µm

A549

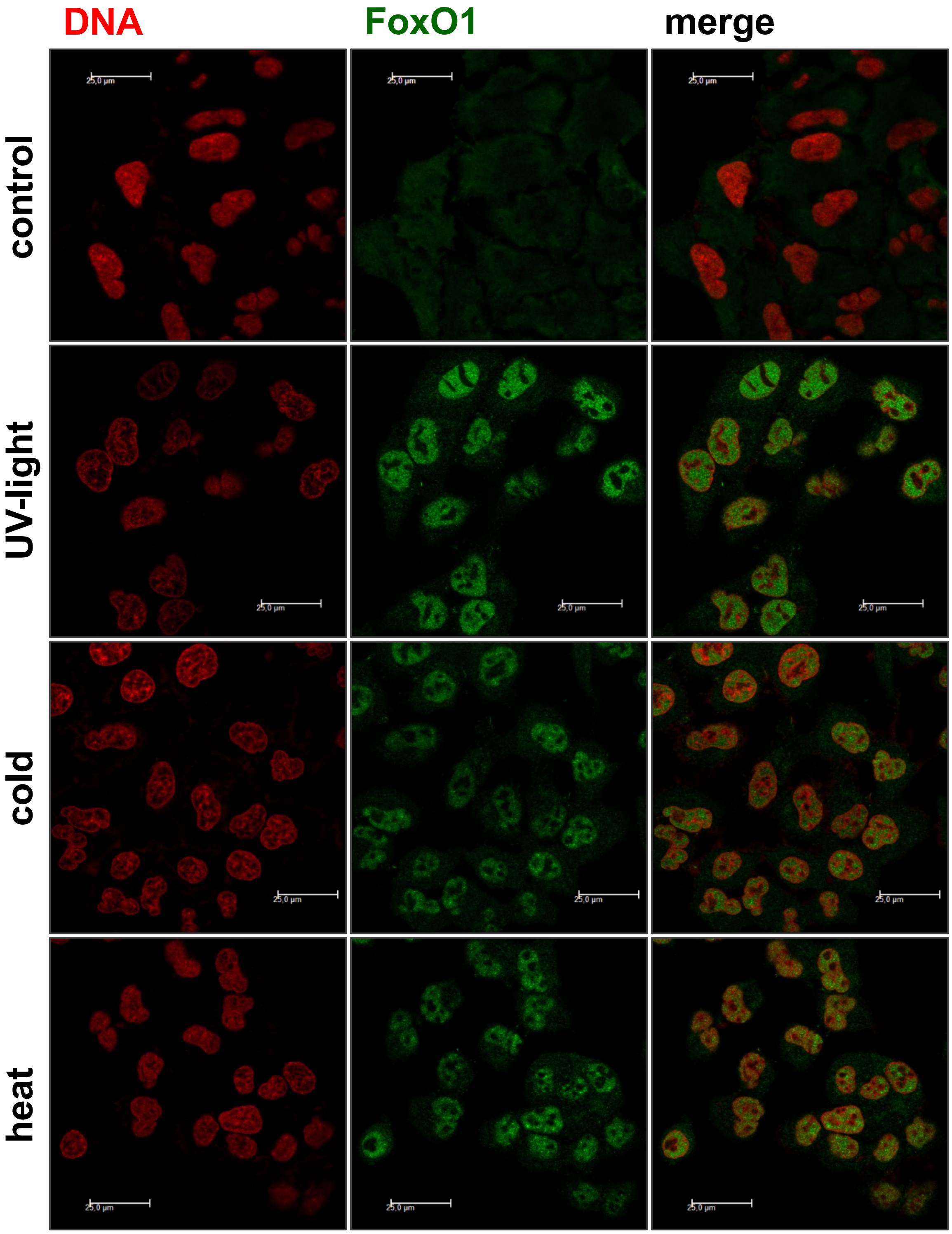

A549

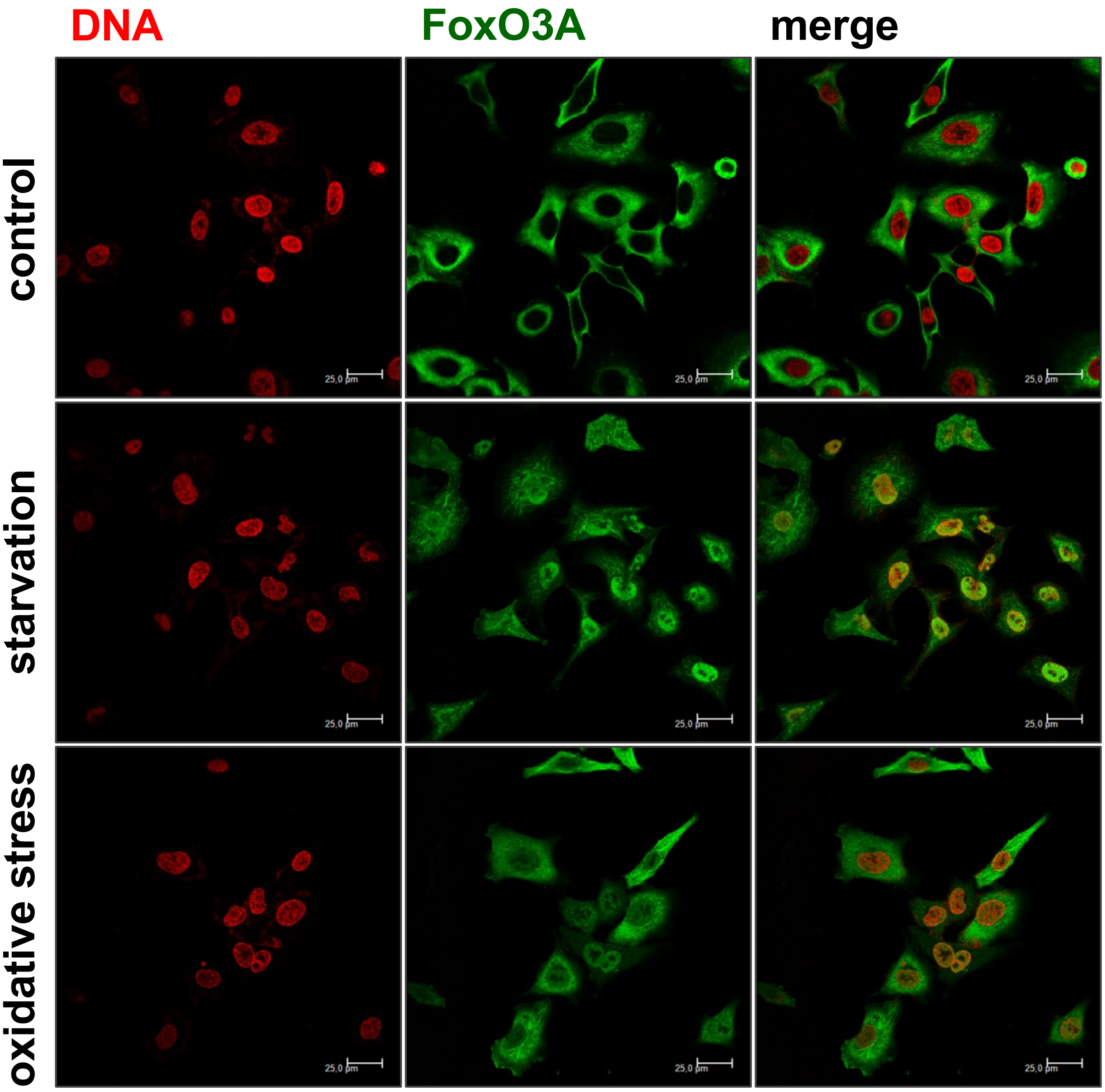

DNA
FoxO3A
merge
control
starvation
oxidative stress
25,0 µm

A549

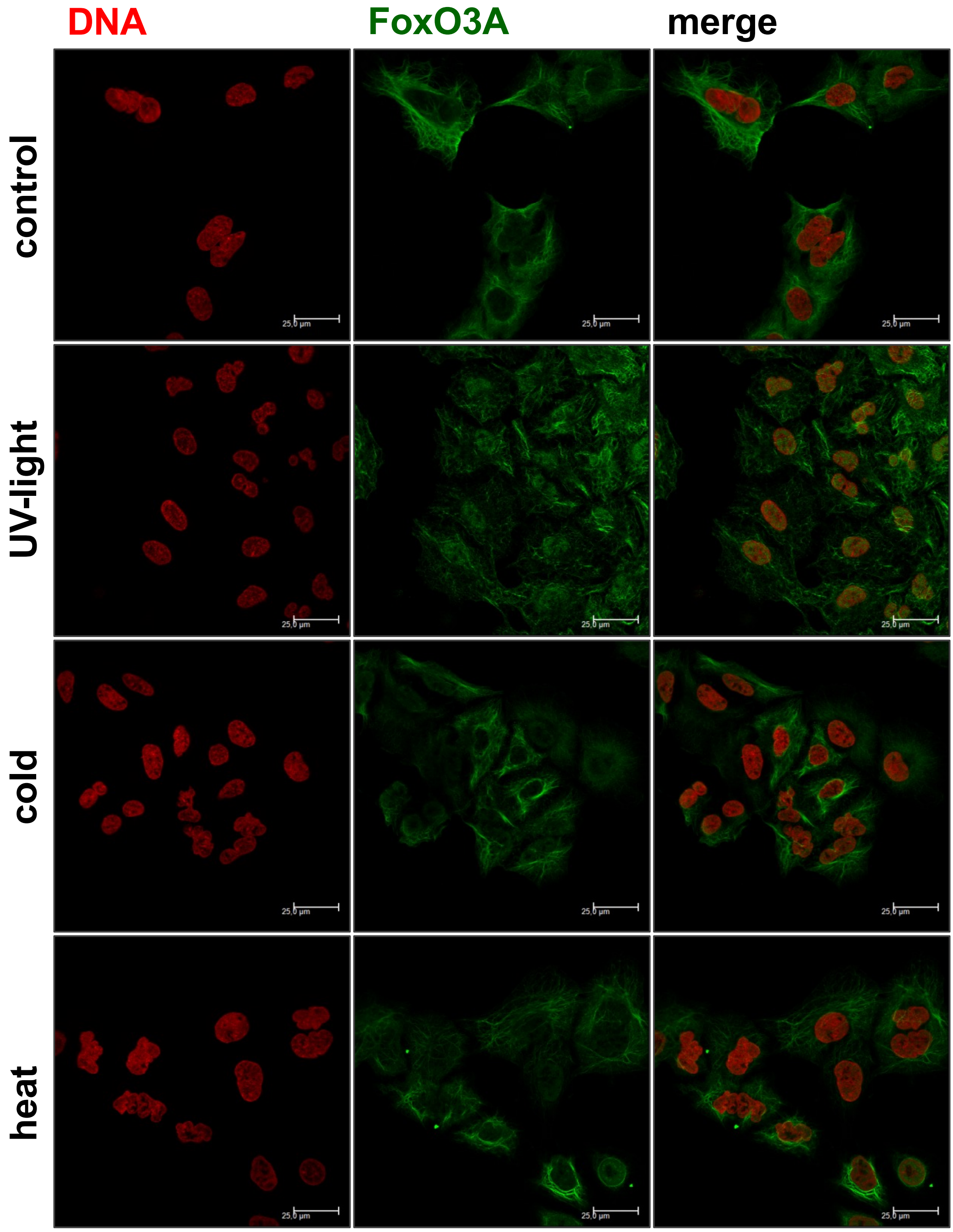

A549

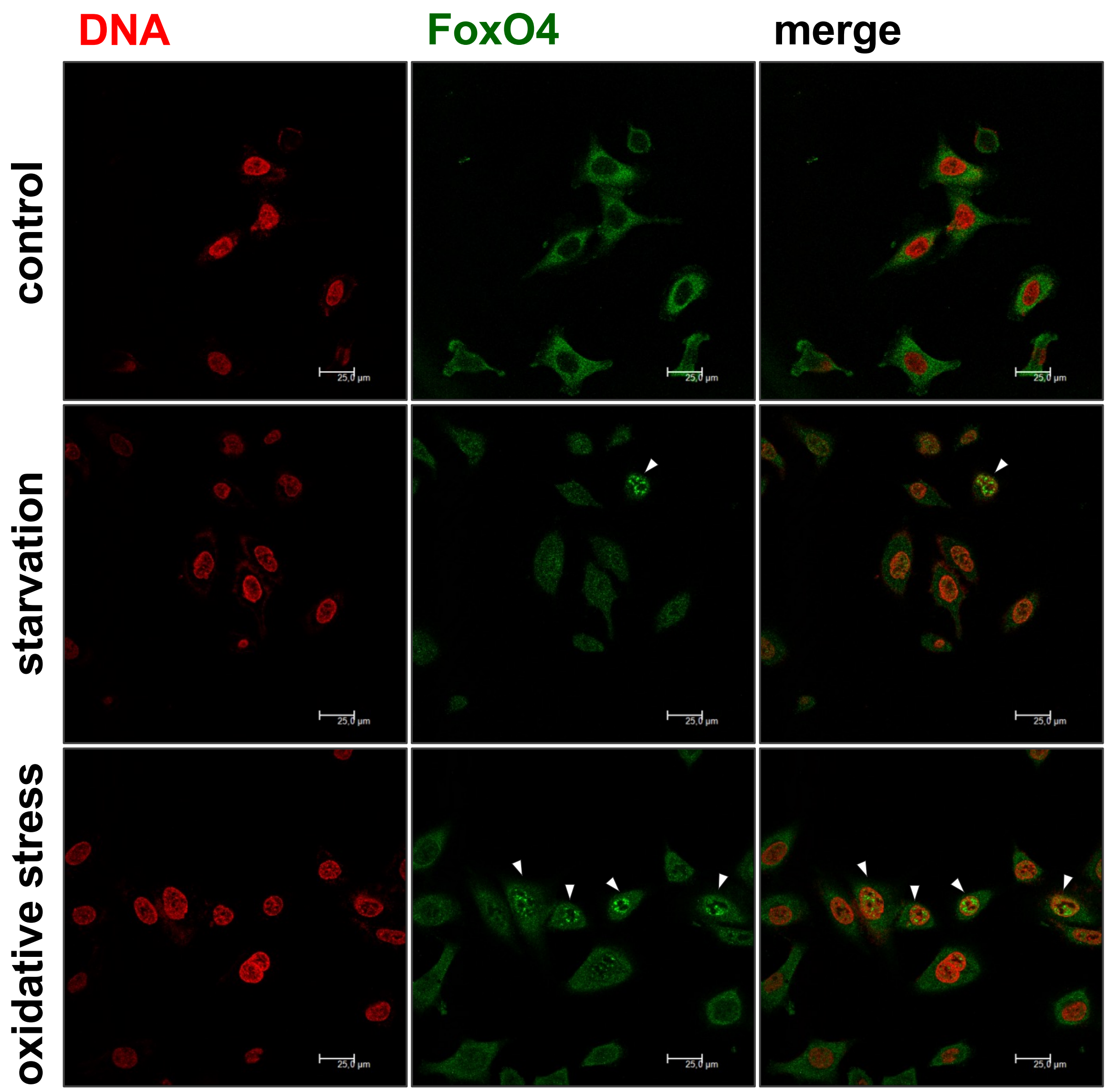
DNA
FoxO4
merge
control
starvation
oxidative stress
25,0 µm

A549

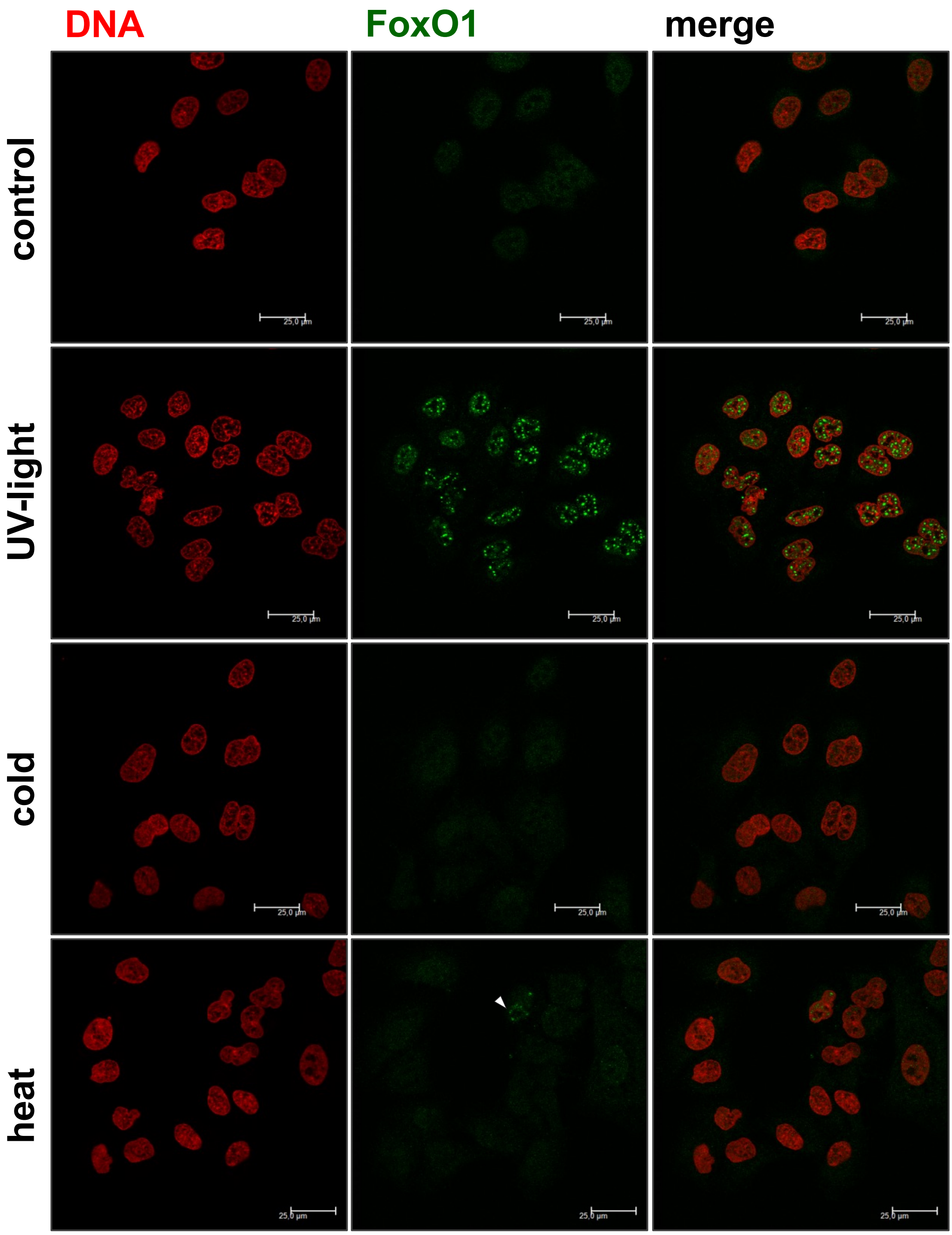

BEAS-2B

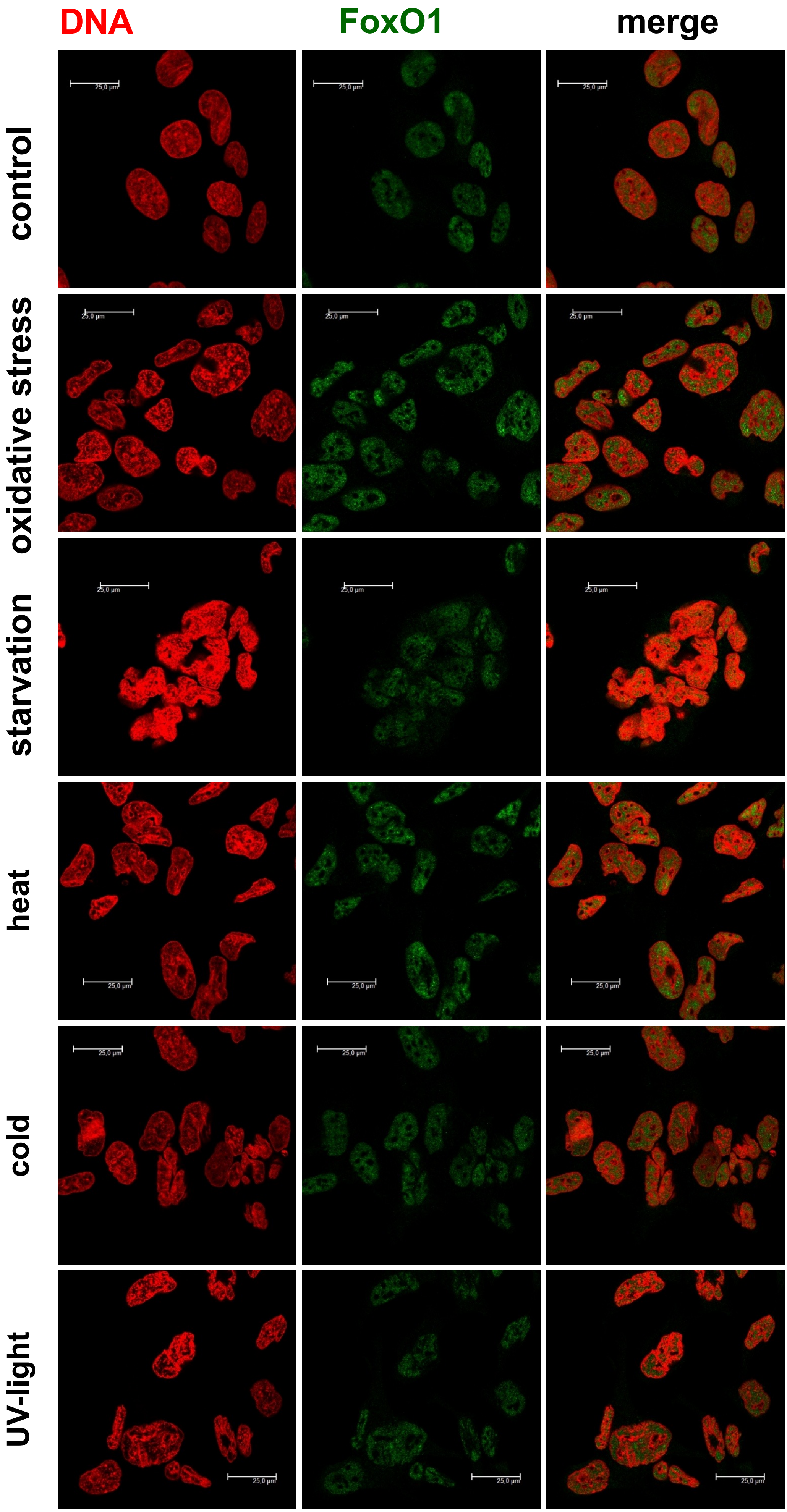

DNA
FoxO1
merge
control
oxidative stress
starvation
heat
cold
UV-light

BEAS-2B

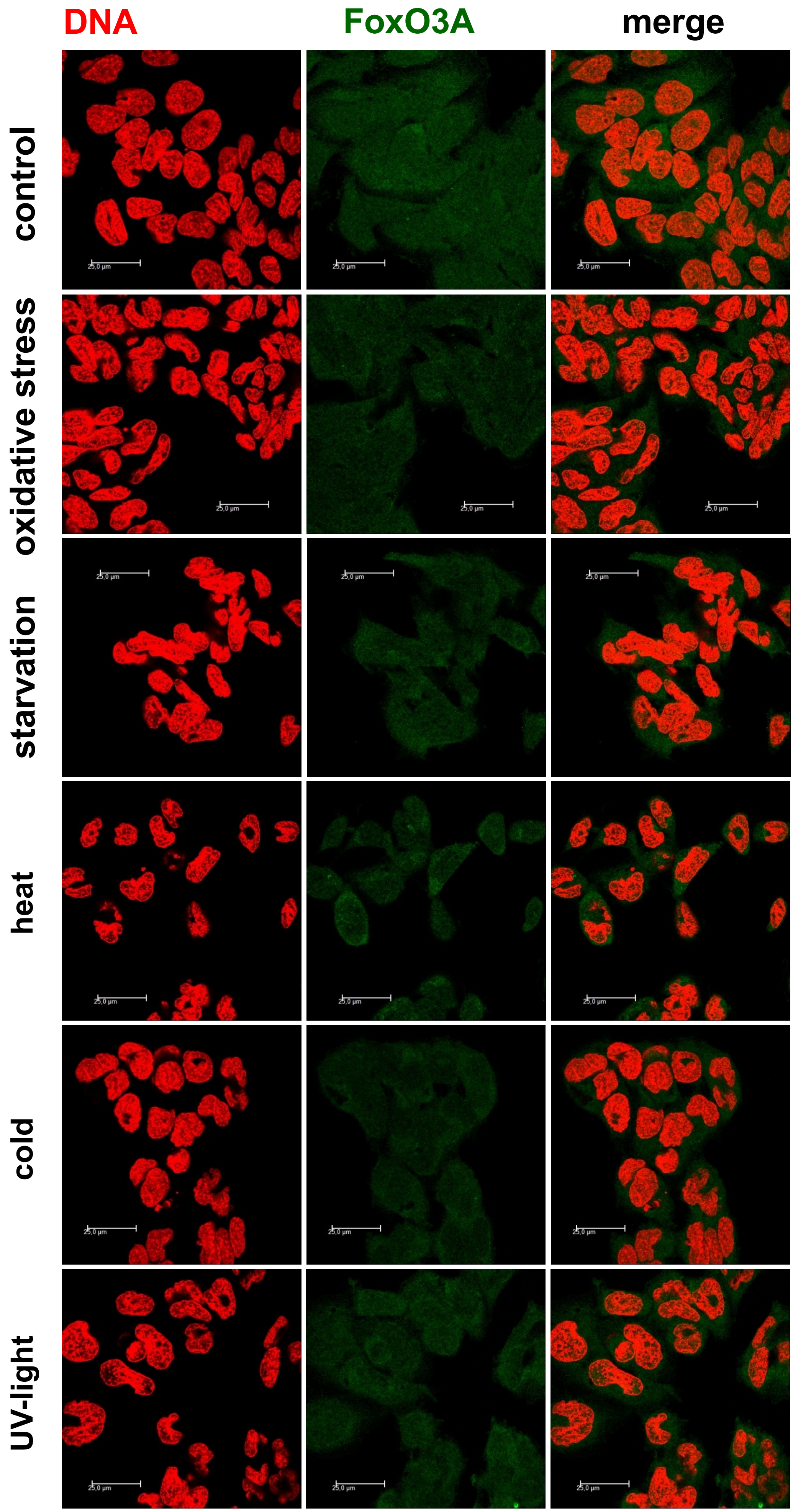

DNA
FoxO3A
merge
control
oxidative stress
starvation
heat
cold
UV-light
25,0 µm

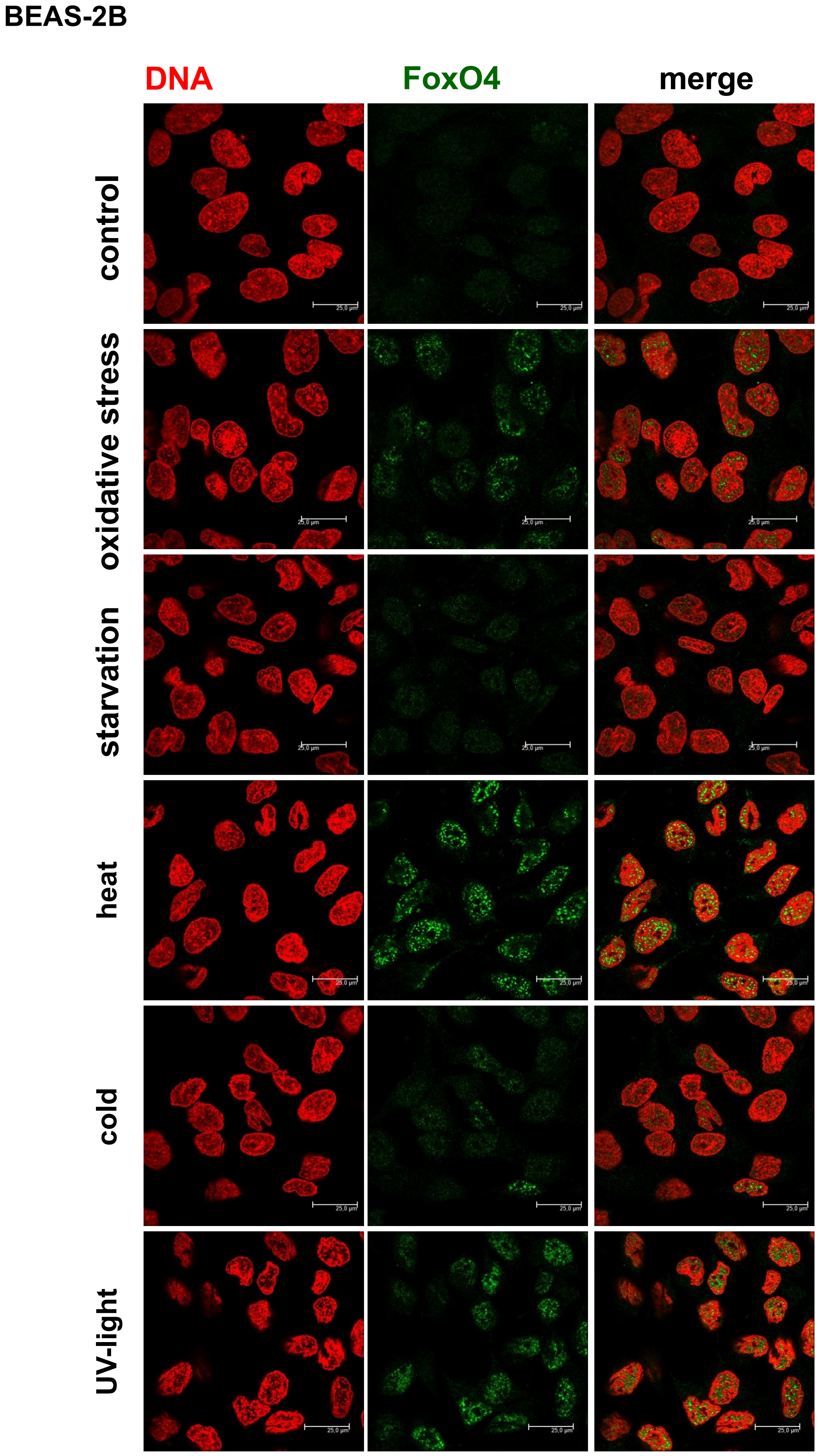

BEAS-2B
DNA
FoxO4
merge
control
oxidative stress
starvation
heat
cold
UV-light
25,0 µm

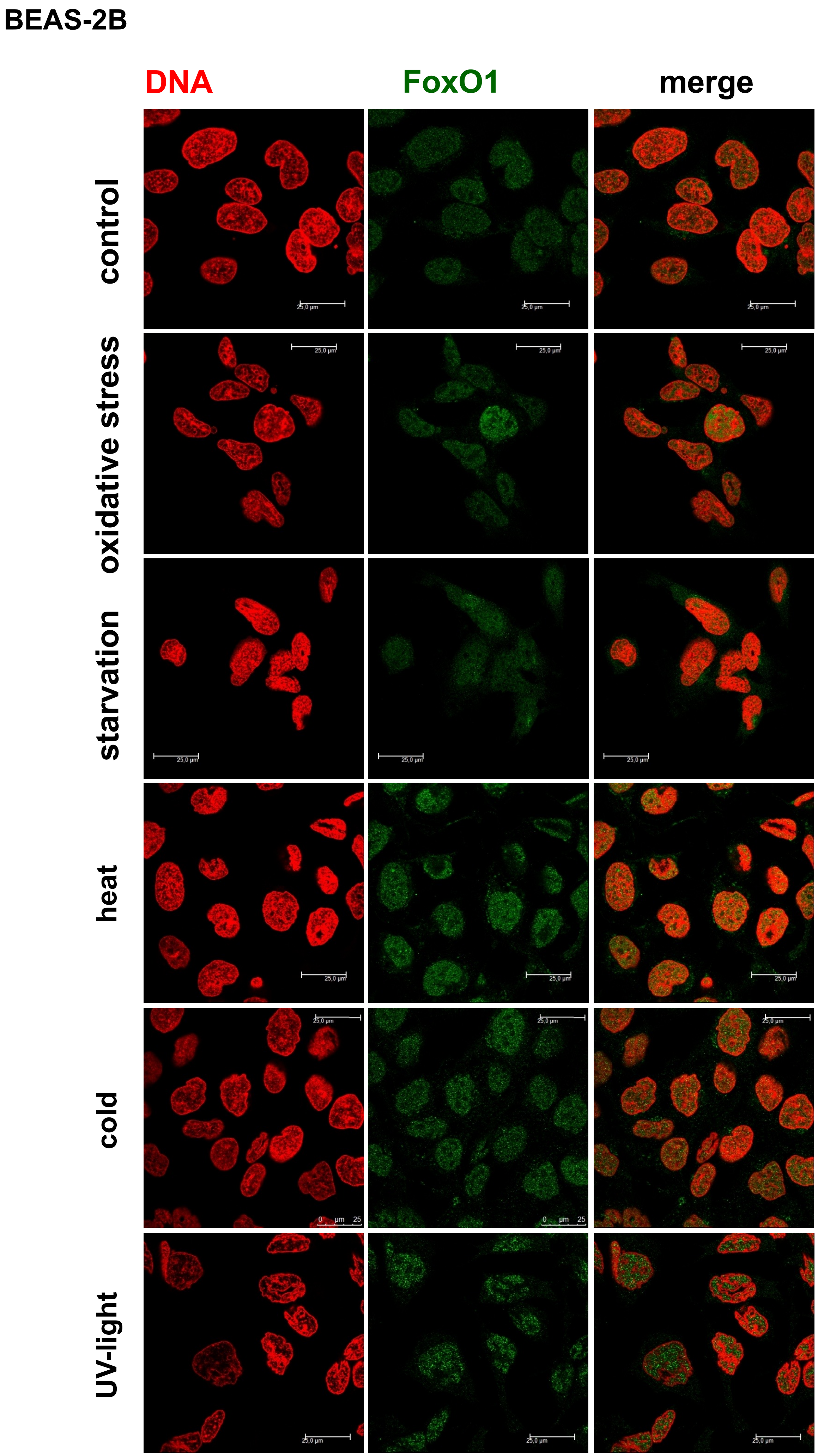
BEAS-2B
DNA
FoxO1
merge
control
oxidative stress
starvation
heat
cold
UV-light
